# From Transcripts to Cells: Dissecting Sensitivity, Signal Contamination, and Specificity in Xenium Spatial Transcriptomics

**DOI:** 10.1101/2025.04.23.649965

**Authors:** Mariia Bilous, Daria Buszta, Jonathan Bac, Senbai Kang, Yixing Dong, Stephanie Tissot, Sylvie Andre, Marina Alexandre-Gaveta, Christel Voize, Solange Peters, Krisztian Homicsko, Raphael Gottardo

## Abstract

Spatial transcriptomics has transformed our ability to map gene expression within intact tissues at cellular and subcellular resolution. Among current platforms, Xenium is widely adopted for its reliability, accessibility, and high data quality. Yet, the properties and limitations of Xenium-derived data remain poorly characterized. Here, we present one of the most comprehensive Xenium datasets to date, encompassing over 40 breast and lung tumor sections profiled using a diverse set of gene panels. Leveraging this resource, we systematically dissect technical noise—including transcript diffusion—alongside assay specificity, panel performance, and segmentation strategies. Our comparison of targeted panels with the newer 5K panel reveals that although the latter captures more transcripts overall, it suffers from reduced per-gene sensitivity and persistent diffusion, even with enhanced chemistry. We demonstrate that single-nucleus RNA-seq (snRNA-seq) markedly improves cell type annotation and enables more precise quantification of diffusion. Building on this, we introduce SPLIT (Spatial Purification of Layered Intracellular Transcripts), a novel method that integrates snRNA-seq with RCTD deconvolution to enhance signal purity. SPLIT effectively resolves mixed transcriptomic signals, improving background correction and cell-type resolution. Together, our findings provide a critical benchmark for Xenium performance and introduce a scalable strategy for signal refinement—advancing the accuracy and utility of spatial transcriptomics.

## Introduction

Recent advances in sequencing- and imaging-based techniques have led to the development of spatially resolved transcriptomics, named Method of the Year in 2020^1^, which enables the spatial quantification of gene expression within tissues. These technologies provide researchers and clinicians with an unprecedented ability to characterize patient samples, including archived FFPE specimens^2^, with high spatial resolution, offering transformative insights that can enhance diagnoses, treatments, and patient outcomes.

In situ hybridization technologies such as Xenium^3^, Merscope^4^, and CosMx^5^ offer high-resolution transcript quantification at the cellular—and even subcellular—level. These platforms—and their earlier iterations—have already generated valuable biological insights and continue to hold significant promise for advancing our understanding of complex tissues^6–8^. However, our understanding of the data characteristics and sources of variability is still maturing as more datasets become available. Critical experimental design questions also remain, particularly around panel selection (e.g., targeted vs. 5K panels for Xenium). While larger panels increase multiplexing capacity, they often come at the cost of reduced sensitivity, higher expense, and longer sample processing times. To date, comparative data between panels remains very limited.

Janesick et al.^3^ were the first to present data generated on the Xenium platform and to perform an in-depth comparison with Visium and Chromium, single-nucleus RNA sequencing (snRNA-seq), using adjacent tissue sections from two samples. Their study demonstrated higher resolution and sensitivity, while also highlighting the potential for integrating multiple platforms. Cook et al.^9^ performed a direct comparison of Xenium and CosMx, using two replicates per platform from a single FFPE block of one patient. Their findings indicated better segmentation in CosMx but higher sensitivity in Xenium. Ren et al.^10^ conducted an in-depth comparison of Xenium 5K, CosMx 6K, and Visium HD but again analyzed only three samples and did not address complex downstream analysis challenges. More recently, Wang et al.^11^ compared Xenium (multiple targeted panels, but no 5K) to Merscope and CosMx but this time using a larger number of samples: seven tumors and sixteen normal tissue types. While they reported similar findings, including higher Xenium sensitivity, their analysis was limited to tissue microarrays (TMA) and focused on specific aspects, without touching on critical factors such as segmentation and transcript diffusion. These studies are for the vast majority, small in scale, typically limited to a few patients at most (unless using TMAs), and none have directly compared targeted Xenium panels to the newer 5K panel in terms of sensitivity and specificity, or transcript diffusion—factors that are essential for robust downstream analyses.

Multiple studies have demonstrated that individual transcripts may be incorrectly assigned to cells, either due to inaccurate segmentation or diffusion into adjacent cells—a phenomenon known as transcript diffusion^12,13^—leading to background contamination that can affect downstream analyses. Transcript misassignments can result in the formation of artificial doublets (or multiplets), where gene expression signatures from different cell types become mixed and ambiguous. While some of this signal mixing can be attributed to diffusion along the z-axis—likely due to overlapping cells causing spillover during x–y plane segmentation—this alone does not fully account for the extent of the contamination, as we explore in this study.

Notably, Mitchel et al. ^14^ demonstrated that misassignment can generate artificial cell-cell correlations (or communication signals), misleading biological interpretations. To address this challenge, they proposed a correction method that leverages single-cell RNA sequencing (scRNA-seq) data and non-negative matrix factorization (NMF) to accurately deconvolve transcript signals and reduce contamination. Ergen and Yosef^13^ framed this problem as transcript diffusion and introduced a probabilistic approach to correct for transcript diffusion. Wu, Beechem, and Danaher^15^ proposed a comparable method for CosMx data, reassigning transcripts to their most likely cell of origin. These findings suggest that transcript diffusion is a widespread challenge across spatial transcriptomic platforms.

We argue that most transcript misassignment arises from diffusion, which can be effectively mitigated through segmentation in transcript space. Importantly, diffusion and segmentation-related misassignment are difficult to disentangle and are often addressed using related computational strategies. For this reason, we refer to both sources of error collectively as diffusion throughout this study. Among available solutions, probabilistic segmentation methods have emerged as a promising approach for correcting diffusion and improving transcript-to-cell assignment accuracy. This approach was first introduced by Petukhov et al.^16^ with their Baysor algorithm. Building on this, Jones et al.^17^ developed ProSeg, a 3D segmentation method that leverages the spatial coordinates of each transcript. ProSeg was specifically designed for emerging spatial transcriptomics technologies and has demonstrated superior performance compared to existing methods, including Baysor and traditional nuclear/cell stain–based segmentation. In addition, several other algorithms have been developed that incorporate transcript-level information, including Segger^18^, BIDCell^19^, and FastReseg^15^, reflecting a broader shift toward data-driven segmentation strategies for resolving transcript diffusion.

While early studies have provided valuable insights and laid the foundation for further advancements, they remain limited in scope, with few publicly available datasets and small sample sizes. Consequently, many findings lack the generalizability required for broader application. Computational methods that leverage transcript-level data have shown clear improvements in signal purity; however, many still depend on heuristic corrections or complex, non-interpretable models, and have been validated on only a limited number of samples. These limitations highlight the need for larger, more representative studies and the development of simpler, more interpretable approaches for correcting spatial transcriptomic data.

Here, we present one of the most comprehensive Xenium datasets generated to date, spanning 41 tissue sections from breast and lung tumors across 27 donors. This resource was designed to investigate key challenges in spatial transcriptomics, including transcript diffusion, assay specificity, panel performance (targeted vs. 5K), segmentation, and cell type annotation. While this study is not intended as an exhaustive benchmarking of existing methods, we share both the dataset and key findings to support the broader community in designing and analyzing spatial transcriptomics experiments. We also demonstrate how a snRNA-seq reference can be used to streamline cell type annotation, evaluate data quality, and mitigate transcript diffusion. To that end, we introduce quantitative metrics for assessing transcript-level data quality, both post-segmentation and following signal correction. Finally, we introduce SPLIT (Spatial Purification of Layered Intracellular Transcripts), a simple yet effective computational method that integrates snRNA-seq with RCTD^20^ deconvolution to enhance signal purity. SPLIT resolves mixed signals caused by transcript diffusion, improves cell-type separation, and is compatible with any segmentation strategy.

## Results

### Data and experimental overview

We generated spatial transcriptomics data from 19 sections derived from formalin-fixed paraffin-embedded (FFPE) blocks of breast cancer tissue and 22 sections from FFPE blocks of non-small cell lung cancer (NSCLC), corresponding to 17 and 10 donors, respectively. Xenium profiling was performed on all samples using the targeted Breast panel for the breast cancer samples, and two targeted, Lung and a custom immuno-oncology panel (Custom IO), and the 5K PRIME panel for NSCLC samples. In addition, matched snRNA-seq and immunochemistry (IHC) data were generated for a selected subset of samples (see Methods and Supplementary Tables 1 and 2 for details).

Segmentation was performed on all Xenium data, resulting in single-cell resolution for subsequent analysis. For samples profiled with targeted panels, we used the default segmentation approach, which applies a 5 µm radius expansion around each nucleus to approximate cell boundaries. In contrast, for samples profiled with the 5K panel, we leveraged the availability of multimodal staining with a 5 µm radius expansion, which combines nuclear, cytoplasmic, and membrane stains to enhance segmentation accuracy.

As described in the next section, snRNA-seq data are used to annotate the Xenium datasets and serve as a reference for evaluating data quality, including gene expression fidelity and cell-type resolution. IHC is used to further validate the annotation—particularly to assess the potential misannotation of specific cell types due to transcript diffusion (see Methods for details).

Figure 1a,b provides an overview of the experimental design, including key metrics such as the number of cells per sample, the number of transcripts per cell, and the number of genes per panel. While we observed substantial variability in cell capture across samples—likely due to differences in tissue size and preservation quality—all samples produced high-quality data, with robust transcript and gene detection at the single-cell level.

**Figure 1.**
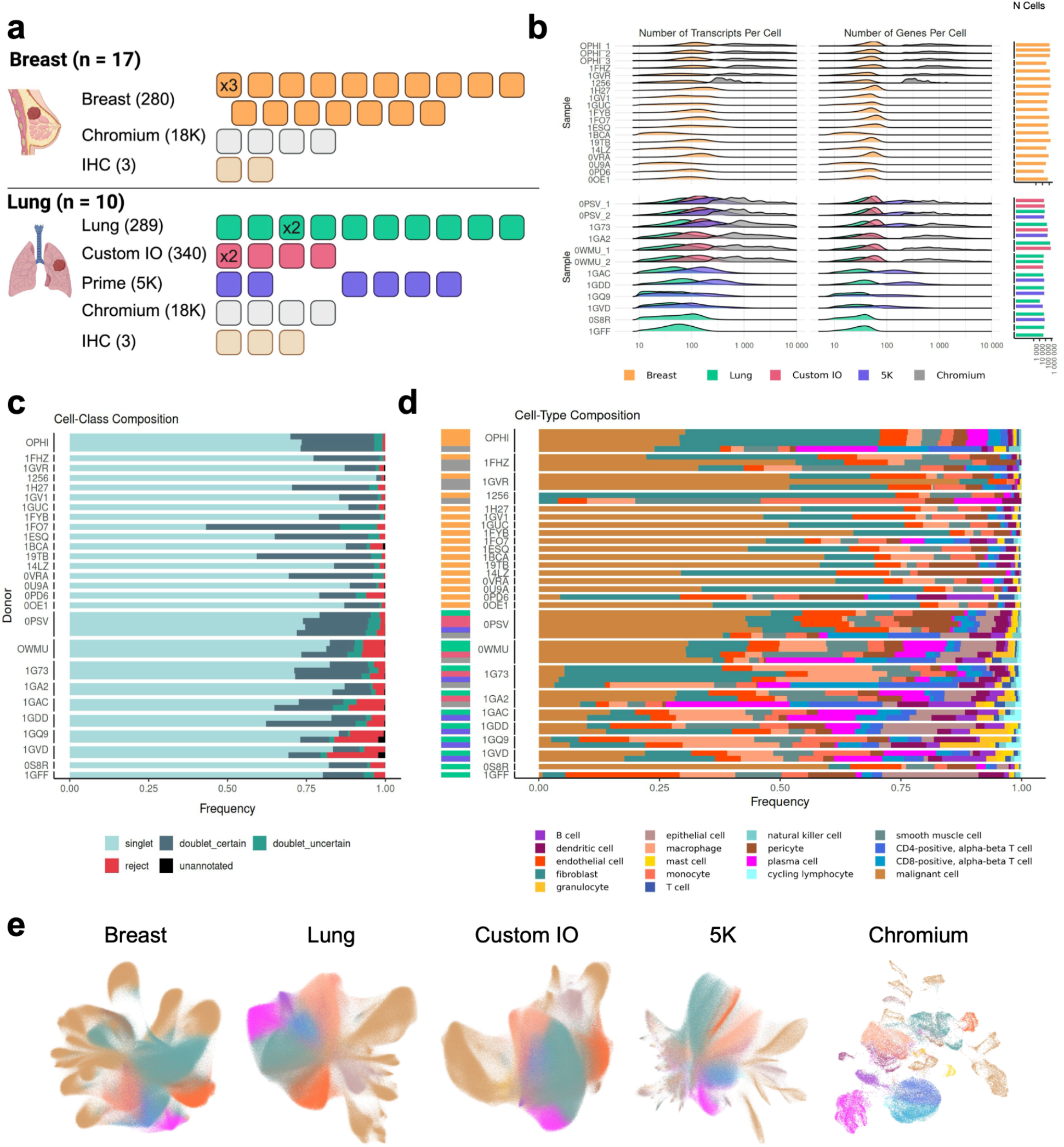
Experimental design and key metrics. **a**, Experiment names correspond to the profiling technology and panel used, with the number in parentheses indicating the panel size. Each sample is represented as a tile, and replicate experiments are marked with “x2” or “x3” within the tile to refer to the number of replicates. Columns group samples originating from the same donor. **b**, Density distributions of the number of transcripts per cell and the number of detected genes per cell for each sample, along with the number of cells per sample, color-coded by panel and assay type. **c**, RCTD spot-class composition for each Xenium sample, grouped by donor. **d**, Cell type composition of all Xenium and Chromium samples, grouped by donor and colored by cell type. The colored strip alongside each row indicates the corresponding panel or assay. Cells in panels c and d were annotated at Level 2.1. **e**, Integrated UMAPs grouped by panel and assay. Cells are colored by cell type as assigned by RCTD using matched Chromium reference at Level 2.1 annotation.

#### snRNA-seq data enable robust annotation of Xenium datasets

While Xenium data inherently provide single-cell resolution, we leveraged matched and annotated snRNA-seq datasets (see Methods) to transfer cell type annotations to each Xenium sample. For this, we applied RCTD^20^, a framework originally developed for spatial transcriptomics deconvolution, which we and others^21^ have found to be highly effective for annotating Xenium data. Importantly, RCTD includes a doublet mode that helps account for mixed signals arising from segmentation errors, overlapping cells, or transcript diffusion (Figure 1c).

Notably, the proportion of cells classified as doublets can serve as a useful quality metric—informing assessments of sample integrity and enabling comparisons across segmentation algorithms. As shown in Figure 1c, Xenium data contain a substantial number of doublets across panels and patients ranging from 1.6% to 54% in breast cancer samples and from 3.5% to 32% in lung cancer samples. Interestingly, the 5K panel appears to have a slightly higher doublet frequency (Supplementary Figure 1a), despite its increased gene content and updated chemistry.

Figure 1d shows the distribution of annotated cell populations across individual samples. The data demonstrate strong reproducibility among technical replicates while revealing substantial variability between patients and between technologies—namely, Chromium and Xenium. This variability is expected: Chromium relies on dissociated single cells and typically requires larger input numbers, a process known to introduce biases in cell-type representation^22^. While snRNA-seq has been shown to mitigate some of these biases, it does not eliminate them entirely^23^. In contrast, Xenium profiles intact tissue sections, preserving spatial context and reducing dissociation-related artifacts.

We also explored the robustness of RCTD in annotating our samples using published (i.e., non-matched) reference scRNA-seq datasets (Supplementary Figure 2). Overall, we found the process to be fairly robust with respect to the reference dataset used, except for some samples where malignant cells were more difficult to annotate (Supplementary Figure 2c). Given the substantial inter-patient heterogeneity in malignant cell transcriptional profiles, our findings highlight the value of a matched reference for cancer research.

Similarly, the annotation remains robust across different Xenium panels, demonstrating that targeted gene panels effectively capture key information for defining major cell types—an aspect we explore further below.

#### Xenium data show great consistency across patients and technical replicates

To evaluate the quality and consistency of Xenium single-cell data, we performed UMAP-based joint analyses across all samples, leveraging the transferred cell type annotations. The data were stratified by panel and assay (Figure 1e). Cells of the same type demonstrated strong concordance across samples without the need for batch correction or additional computational integration, aside from Seurat log normalisation. This indicates that both Xenium and Chromium platforms introduce minimal technical variation. Supplementary Figures 3 and 4 present the same dataset faceted by sample, revealing that non-malignant cell populations align consistently across individuals, whereas malignant cells cluster in a patient-specific manner, as anticipated. Moreover, data from technical replicates derived from adjacent tissue sections show near-perfect concordance (Supplementary Figure 5a), confirming the high reproducibility of the platform.

#### Targeted gene panels outperform the 5K panel in sensitivity

As expected, Figure 1 shows that the 5K panel detects a higher total number of transcripts and genes. However, this increase does not translate into improved cell-type separation in UMAP space (Figure 1e). To investigate this further, we focused on genes shared across panels (n = 194) to assess whether the 5K panel exhibits reduced sensitivity when controlling for gene content.

Figure 2a shows that the 5K and Lung panels yield a comparable representation of cells in the UMAP space when analysis is restricted to shared genes. However, approximately 60% of cells in the 5K dataset fail quality control due to low transcript counts, indicating reduced sensitivity compared to the Lung panel. This discrepancy in data retention points toward underlying sensitivity differences between the panels. Supporting this, Figure 2b shows that, for the same subset of common genes, the 5K panel detects significantly fewer transcripts than the targeted Lung panel. This observed reduction in sensitivity is consistent with known limitations of the 5K panel and likely reflects protocol modifications like the reduced probe count to limit optical overcrowding.

**Figure 2.**
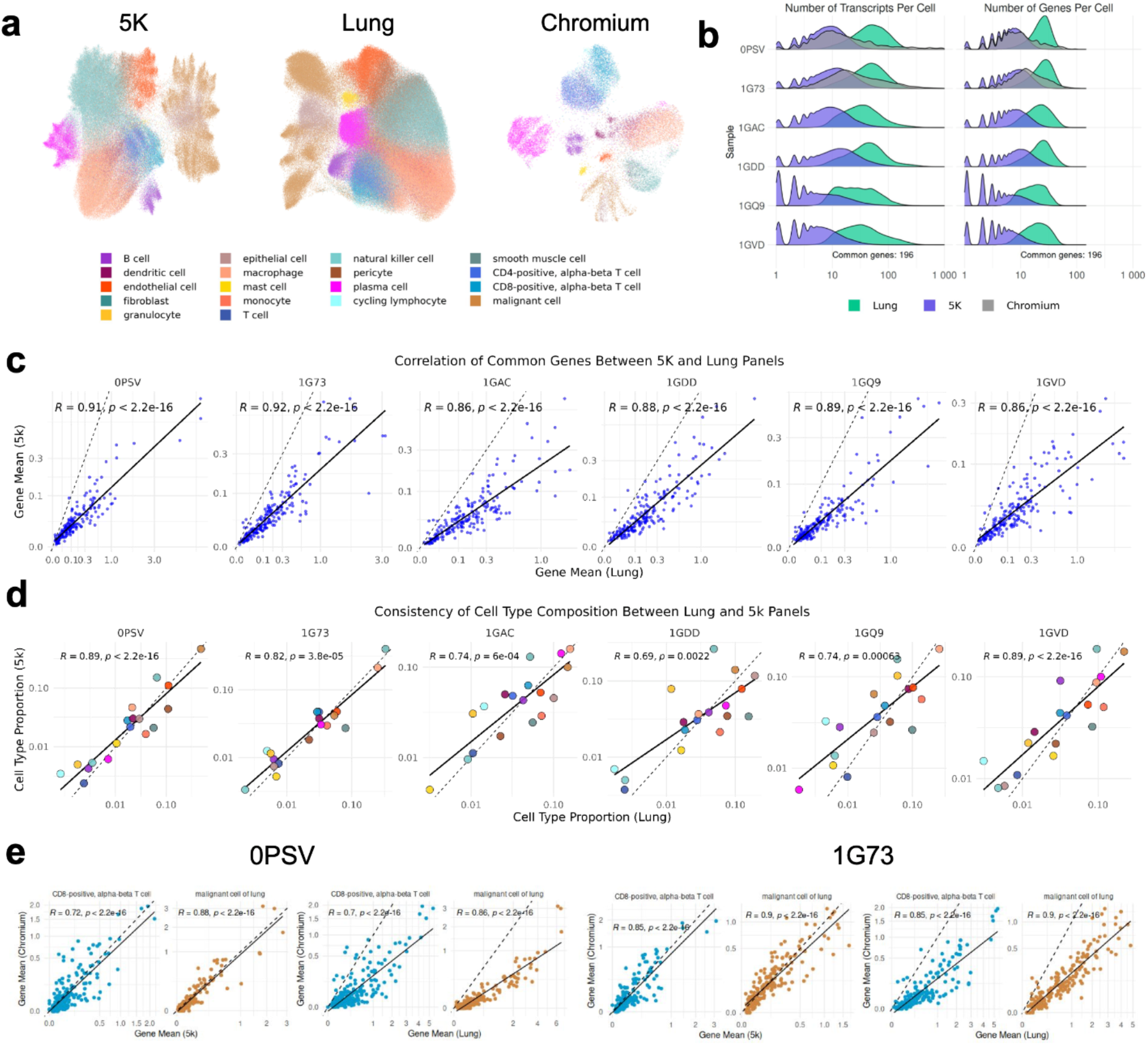
Comparison of Xenium prime (5K) and targeted (Lung) panels. **a**, UMAP visualization of lung cancer samples based on 194 genes common to the Xenium 5K panel, Xenium Lung panel, and Chromium data. Cells are colored according to their annotated cell types. **b**, Distribution of transcript counts per cell in the Xenium 5K, Xenium Lung, and Chromium datasets, restricted to the common genes. **c**, Scatterplot showing average expression of common genes in matched samples from the Xenium 5K and Xenium Lung panels. **d**, Scatterplot showing cell type composition in matched samples from the Xenium 5K and Xenium Lung panels. **e**, Scatterplots comparing average expression of the 194 common genes between Chromium and Xenium Lung, and between Chromium and Xenium 5K, in CD8^+^ T cells and malignant cells. Analyses are based on matched samples from two donors (0PSV and 1G73) profiled with all three assays. Extended comparisons across all cell types, and a direct comparison between Xenium Lung and Xenium 5K panels, are shown in Supplementary Figures 5 and 6.

We also assessed gene expression consistency between adjacent tissue sections using shared genes across panels. As shown in Figure 2c, the 5K and targeted panels exhibit a reasonable correlation, though the 5K panel consistently shows lower sensitivity. Notably, the 5K sensitivity is comparable to Chromium (Figure 2b), underscoring a key trade-off between gene breadth and detection depth. While the 5K panel covers a broader set of genes, these results highlight important limitations in sensitivity. Whether this expanded target list compensates for the reduced per-gene detection remains an open question.

#### Targeted and 5K panels show high concordance in cell-type compositio

Despite the reduced sensitivity of the 5K panel, we observe good consistency in cell type composition between adjacent sections profiled with different panels (Figure 2d), suggesting that the broader gene coverage of the 5K panel might help compensate for its reduced sensitivity. Similar results are observed when comparing the 5K panel to the Custom IO targeted panel (Supplementary Figure 5e). Interestingly, the consistency of cell type proportions is highest between the two targeted panels—Custom IO and Lung—even though they share a relatively small number of genes (n = 87) (Supplementary Figure 5g). This suggests that Xenium is robust to variations in gene panel design—at least within the context of targeted panels that share the same underlying chemistry. To test whether better agreement between targeted panels is explained by segmentation differences, we reanalyzed 5K data using the same method but found no significant improvement in correlation (Supplementary Figure 5d,f).

To further evaluate the similarity of cell type profiles between the Xenium platform (5K and targeted panels) and the Chromium reference, we computed correlations across matched donor samples. As shown in Figure 2e and Supplementary Figure 6, both Xenium panels exhibit good agreement with Chromium-derived profiles, with the 5K panel resembling the Chromium data more in terms of sensitivity.

#### Targeted gene panels provide extensive information for defining major cell types

To further evaluate the effect of gene set size on cell-type resolution, we reanalyzed Chromium data using only the genes included in the targeted panels. As shown in Supplementary Figure 5h, Chromium data achieves a clear separation of cell types despite the reduced gene sets. Moreover, this separation persisted even when the analysis was further constrained to a subset of 194 genes shared between the targeted Lung and 5K panels (Figure 2a). These findings suggest that the diminished resolution observed in the Xenium data is not solely due to a limited number of genes analyzed.

Notably, even after restricting to shared genes, the Lung and 5K panels still failed to achieve the same level of cell-type separation as observed with Chromium data. This supports the idea that transcript diffusion between neighboring cells may blur cell-type boundaries, limiting resolution in spatial transcriptomics.

#### Xenium data exhibit substantial transcript diffusion

In Xenium data, transcripts are assigned to cells based on a computational tissue expansion radius, which extends cell boundaries by 5 μm beyond the segmented nucleus, or until another cell boundary is reached. This default setting, defined by 10x genomics, aims to capture the mRNA dispersed throughout the cell cytoplasm. Based on our RCTD analyses (see Methods), we find that this expansion often results in the assignment of mixed transcriptional signals to individual cells, likely representing “doublets”. Supporting this, the doublet rate is lower when no tissue expansion is applied (0 μm; Supplementary Figure 7a), suggesting that these admixtures are due to segmentation errors or RNA diffusion between neighboring cells. While some studies attribute such admixtures primarily to vertical (z axis) overlap^24^, others show frequent signal mixing within the same z-plane ^14^. We therefore also investigated the contribution of horizontal transcript diffusion to signal contamination.

To test this, we examined the correlation between the extent of contamination — estimated by the weight of the secondary cell type signal in a cell — and the abundance of that same cell type in the local 2D neighborhood (Figure 3a,b), providing a local measure of diffusion potential. As shown in Figure 3c, these two metrics are strongly correlated, suggesting that higher local concentrations of a given cell type increase the likelihood of contamination. Additionally, cells classified as doublets have a larger abundance of the secondary cell type in the neighborhood than singlets (Figure 3d and Supplementary Figure 7b), further supporting this hypothesis.

**Figure 3.**
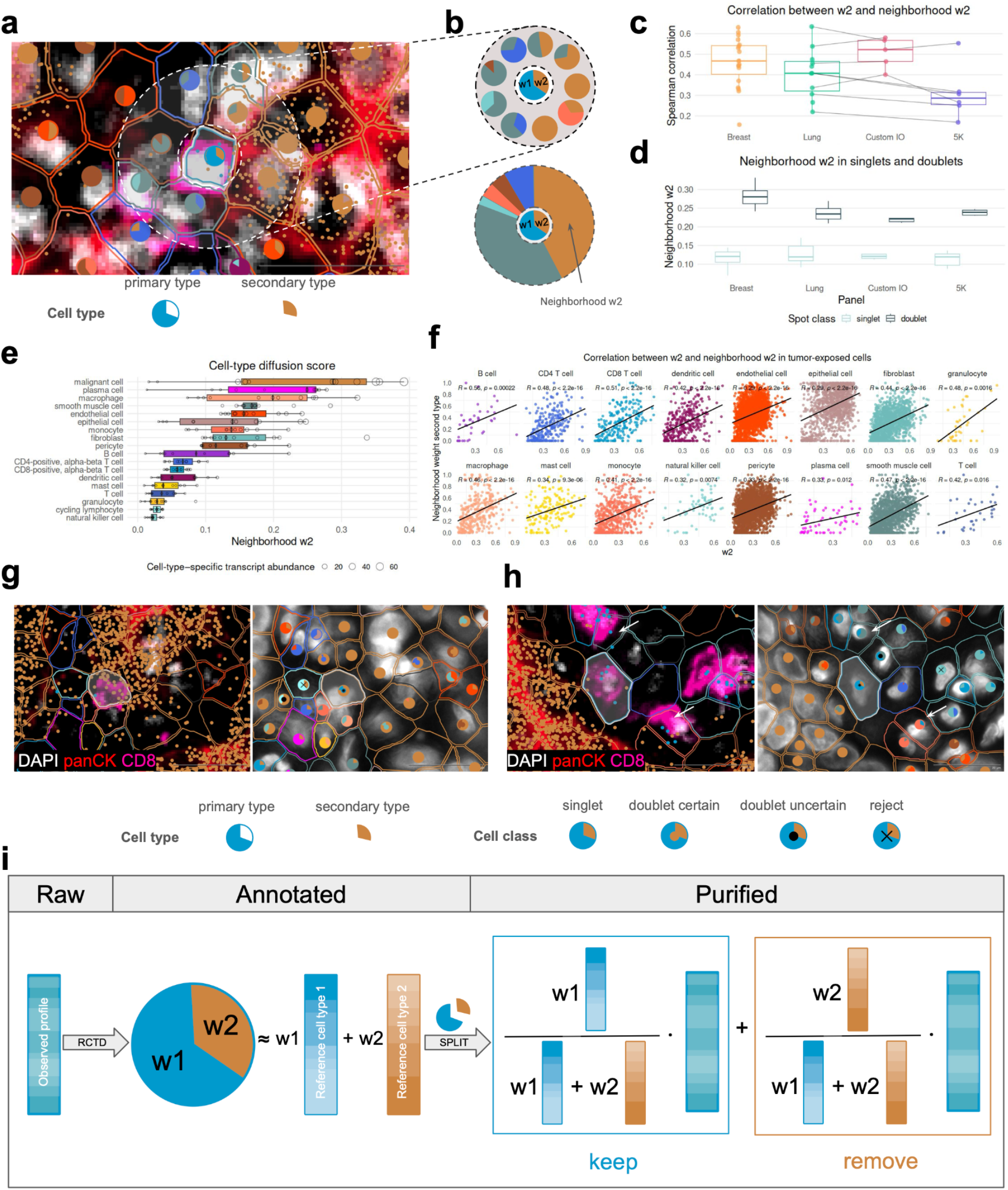
Transcript diffusion in Xenium data and the SPLIT purification approach. **a**, Section of a Xenium sample showing cell segmentation overlaid with RCTD-derived cell type annotations. Border colors indicate the primary cell type assigned to each cell. Pie charts represent RCTD-computed cell type compositions, where sector size corresponds to the proportion of each cell type and color denotes identity. Small dots represent transcripts specific to tumor cells (brown) and CD8^+^ T cells (blue). **b**, Schematic illustrating the definition of a cell’s neighborhood (top) and its neighborhood composition (bottom). The proportion of the cell’s secondary cell type (brown) within its spatial neighborhood is defined as the neighborhood w2. Colors represent distinct cell types. **c**, Spearman correlation between a cell’s secondary cell-type weight (w2) and the average proportion of the same cell type in its neighborhood (neighborhood w2), across all cells and samples. Each dot represents one Xenium sample; lines connect samples from the same donor. **d**, Distribution of mean neighborhood w2 in cells classified as singlets or doublets by RCTD. **e**, Cell-type diffusion score, defined as the average neighborhood w2 per secondary cell type shown for all Xenium samples profiled with the Lung panel (see Supplementary Figure 8 for other panels). Each circle represents one sample; the circle size reflects cell type transcript abundance (see Methods). **f**, Spearman correlation between w2 and neighborhood w2 in tumor-exposed cells (cells where the secondary cell type is “malignant cell”), faceted by primary cell type. **g**, IHC and morphology image showing a cell identified as a CD8^+^ T cell by IHC but annotated as a tumor cell by RCTD due to high levels of tumor-specific transcripts. The cell is highlighted with an asterisk (*). The border color indicates the RCTD-assigned primary cell type. Dots represent cell-type–specific transcripts (tumor in brown, CD8^+^ T cell in blue). **h**, IHC and morphology image of a cell confidently annotated as a CD8^+^ T cell by both IHC and RCTD (singlet), displaying residual tumor-specific transcripts. Arrows indicate cells where the primary and secondary labels appear to be swapped. The border color represents the RCTD-assigned primary cell type; dots show tumor (brown) and CD8^+^ T cell (blue) transcripts. **i**, Schematic of the RCTD cell type annotation in doublet mode. Each cell is modeled as a mixture of two cell types (primary and secondary) with proportions w1 and w2. SPLIT decomposes this mixture into purified primary and secondary components based on reference profiles and assigned weights. By default, the primary profile is retained and used in downstream analysis. Panels a, f, g, and h correspond to samples from the same donor (Lung 0PSV).

Notably, the strength of the diffusion potential varies by cell type, with the most pronounced effects observed in cell types with higher RNA abundance (Figure 3e and Supplementary Figure 8). This trend is intuitive, as cells with greater RNA content and that are more abundant have greater diffusion potential. Since malignant cells exhibit the highest diffusion potential, we selected them to illustrate how tumor-derived signals contaminate other cell types. Figure 3f demonstrates a clear correlation between the local abundance of malignant signals and the extent of contamination in neighboring cells.

To further illustrate this effect and provide orthogonal validation, we performed multiplexed IHC on a subset of samples following Xenium analysis (see Methods). Although IHC has its limitations, protein staining is generally regarded as more robust and less prone to diffusion artifacts, offering an independent modality to validate spatial transcriptomic data and cell type annotations. In particular, the IHC data allowed us to confidently identify CD8^+^ T cells and malignant cells. Consistent with our hypothesis of physical transcript diffusion, Figure 3a,g,h shows tumor-specific transcripts extending into adjacent CD8^+^ T cells, supporting the presence of diffusion-driven contamination. In some instances, this contamination is sufficiently strong that RCTD confuses the primary and secondary cell types (Fig. 3g). Notably, even in cells classified as singlets by RCTD, residual expression of transcripts from neighboring cell types is detected (Fig. 3h), indicating that contamination can occur even in confidently annotated single-cell profiles.

### Signal decomposition reduces diffusion and enhances cell type purity

As shown in Figures 1 and 3, Xenium data exhibit mixed signals due to transcript diffusion. Supplementary Figure 1b demonstrates that removing RCTD-identified doublets and focusing only on singlets improves visual separation between cell types but significant blurring between populations remains. While one might consider simply discarding doublets as a straightforward solution, this approach is unsatisfactory for two main reasons: (1) it would exclude a substantial fraction of cells — and their associated transcripts — from analysis, and (2) even cells classified as singlets can still exhibit mixed signals, though not enough to be flagged as doublets. This is evident in Figure 3h, where singlet-classified cells still show signs of contamination.

To address this challenge more effectively, we propose a refined strategy to correct for spatial diffusion by decomposing the signal within each cell showing signs of contamination (see Methods for details). Our approach leverages RCTD and cellular decomposition to separate mixed signals into cell type–specific components, thereby minimizing contributions from secondary cell types. Specifically, RCTD is used to model each cell’s expression profile as a weighted combination of reference cell type profiles (Figure 3i). These weights are then used to proportionally assign transcripts to their most likely sources—primary or secondary—based on cell type decomposition (see Methods).

Using this strategy, we introduce a new framework called SPLIT (Spatial Purification of Layered Intracellular Transcripts), which separates mixed signals by splitting each cell according to its inferred cellular composition. By default, SPLIT decomposes all cells — both singlets and doublets — that show evidence of a secondary signal, by preserving the expression profile of the primary cell type (Figure 3i).

A UMAP embedding of the resulting SPLIT-corrected cells (Supplementary Figure 9a) reveals more distinct clustering and improved separation compared to the original mixed signals. SPLIT offers the option to limit correction to only doublets. Additionally, SPLIT can leverage spatial information to assess the abundance of secondary signals in the local neighborhood (i.e., local diffusion potential), enabling selective decomposition only when contamination is likely. This spatially informed strategy helps prevent overcorrection of phenotypes that may be underrepresented or absent in the reference (see Discussion). For instance, cycling tumor cells—missing from our reference but present in some matching Xenium samples—were often classified as doublets with malignant cells as primary and cycling lymphocytes as secondary type. SPLIT avoids decomposing such cells, preserving distinct and biologically relevant populations (Supplementary Figure 10).

Beyond background correction, SPLIT also enables phenotype refinement by shifting the primary phenotype assignment to one of the secondary phenotypes, based on transcriptional neighborhood homogeneity. We refer to this process as *SPLIT-shift* (Supplementary Figure 9b).

#### SPLIT compares favorably to other correction methods

Next, we evaluated SPLIT’s performance relative to other correction methods, namely ResolVI^13^ and ovrlpy^24^ as well as to various segmentation approaches. Some of these segmentation methods (i.e., ProSeg) also directly model transcript diffusion. Since SPLIT and other signal enhancement techniques can be combined with any segmentation method, we assessed all combinations in our comparison.

In visual assessments, SPLIT led to clearer separation of cell types in UMAP space (Figure 4a). To further evaluate the performance of correction methods, we compared each method across different segmentation outputs based on cell type separation, batch integration, and the retention of cells and transcripts. An optimal correction method should enhance separation between cell types, avoid introducing or amplifying batch effects, and preserve as many cells and transcripts as possible.

**Figure 4.**
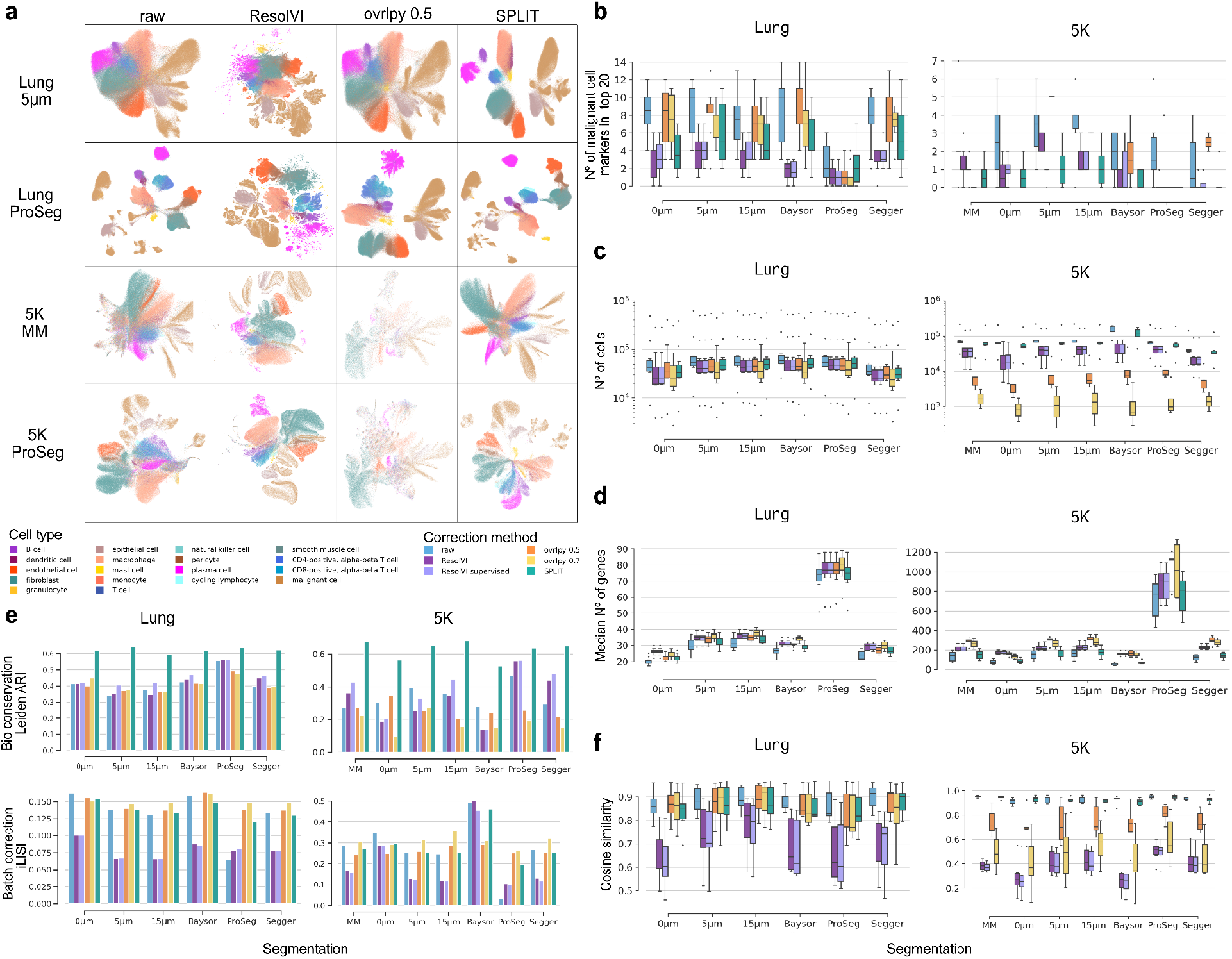
SPLIT improvements and comparison of segmentation approaches. **a**, UMAPs of Xenium cell-level data obtained before (raw) and after count correction for Lung and 5K panel, and default (i.e., 10x 5μm for Lung panel and 10x mm 5μm for 5K panel) or ProSeg segmentation. **b**, Strength of T cell contamination by malignant cells. Assessed by the number of malignant marker genes (with N = 20 markers) found in the top 20 ranks of sorted logistic regression coefficients, obtained before (raw) and after count correction for different segmentations and count correction methods. **c**, Number of cells obtained before (raw) and after count correction for different segmentations and count correction methods. **d**, Median number of genes obtained before (raw) and after count correction for different segmentations and count correction methods. **e**, Biological conservation and batch correction metrics from SCIB (higher is better) for Lung and 5K panels. **f**, Cosine similarity of Xenium with Chromium snRNA-seq T cell pseudo-bulk profiles obtained before (raw) and after count correction for different segmentations and count correction methods.

Additionally, to evaluate transcript diffusion, we adopted an approach introduced by Mitchel et al.^14^. In this method, cells of a given cell type *i* (e.g., T cell) are assigned a binary label indicating whether they are distant (0) or adjacent (1) to a cell of a contaminating cell type *j* (e.g., malignant cell). A logistic regression model is trained to predict the presence of cell type *j* in the neighborhood based on the expression profiles of cell type *i*. Ideally, the predictive genes should not be specific to the contaminating cell type; if they are, it suggests that transcript diffusion is contributing to the observed signal.

Our analysis focused on malignant cells and T cells: malignant cells are highly transcriptionally active and tend to release large amounts of RNA into their surroundings, while T cells are smaller, contain fewer transcripts, and are thus more vulnerable to contamination (Figure 3e-h). Additionally, we had access to IHC data for both phenotypes, which enabled visual inspection of the results.

Overall, all correction methods improve cell type separation and decrease contamination compared to raw segmentation output, across all segmentation methods. The greatest separation and lowest contamination are observed when combining ProSeg with count correction, even though ProSeg already incorporates a correction for transcript diffusion in its probabilistic model (Figure 4b, 4e top row). We further observe that 10x nucleus segmentation, sometimes assumed to be a “ground truth” or free of contamination ^17^, still shows signs of contamination which are improved by count correction (Figure 4b). On the other hand, samples run with the 5K panel displayed much lower contamination compared to other gene panels, especially when using multimodal segmentation (Figure 4b). However, this reduction may be attributed to decreased statistical power resulting from a lower number of transcripts detected per cell. Batch effects are overall unchanged by correction methods, except for ResolVI which sometimes results in slightly worse iLISI (Figure 4e, bottom row).

SPLIT results in the best cell type separation (Figure 4a,e top row) and cosine similarity with Chromium snRNA-seq (Figure 4f), while ovrlpy and ResolVI remove more of the contaminating signal (Figure 4b). However, ovrlpy and ResolVI result in a substantial decrease in the number of genes expressed, resulting in many empty or near-empty cells. Thus the number of cells decreases significantly after reapplying QC thresholds (Figure 4c). Once these low-expression cells are removed, the remaining cells express a higher median number of genes compared to raw segmentation data (Figure 4d). ProSeg yields a higher median gene count, though it includes many low-expression values. This is likely due to its probabilistic model, which assigns non-zero expression levels to genes that may otherwise be undetected. Our data further demonstrate that segmentation algorithms can markedly impact cell-level transcript quantification. As shown in Supplementary Figure 7a, newer methods—particularly ProSeg—significantly reduce the frequency of RCTD-assigned doublets and enhance cell-type separation, leading to improved accuracy in downstream analyses.

## Discussion

In this study, we leveraged an extensive Xenium spatial transcriptomics dataset to assess data quality, evaluate segmentation and correction strategies, and introduce SPLIT, a novel method for signal purification. Our analyses confirm that Xenium produces high-quality, highly reproducible data with low technical variation. These characteristics enable reliable integration across tissue sections and even across patients, offering strong potential for both single-sample and multi-sample spatial studies.

Our comparison of panel designs highlights that targeted panels, despite covering fewer genes, offer greater sensitivity and are sufficient to resolve major cell types. In contrast, the 5K panel, though more comprehensive in coverage, exhibits reduced per-gene sensitivity and poorer performance in downstream analyses. This trade-off suggests opportunities to optimize probe composition by fine-tuning panel content or supplementing the 5K panel with additional targets. This is something that is recommended by 10x genomics, and recent computational work in probe design and selection could also be used ^25^. As spatial transcriptomics scales to larger discovery projects, it will be important to systematically quantify this sensitivity-versus-coverage trade-off and establish criteria for optimal panel design. Depending on the goals of the study, particularly in large-scale, exploratory settings, this trade-off may ultimately be acceptable.

We also confirmed previous reports that single-cell expression data derived from Xenium are prone to transcript contamination, stemming from segmentation inaccuracies and transcript diffusion—two interrelated sources of error. Using complementary RCTD deconvolution and IHC data, we showed that this artifact is widespread and disproportionately affects low RNA-content cells, such as T cells. As noted by Mitchel et al.^14^, and further validated in our analysis, such contamination can significantly distort the inference of cellular niches—one of the key goals of spatial transcriptomics. These findings underscore the importance of developing robust computational strategies to correct for contamination and highlight the need for researchers to critically evaluate the outputs of cell–cell interaction inference tools. This concern is particularly relevant for emerging foundation models trained on large-scale spatial transcriptomics datasets, where self-supervised tasks often involve predicting masked cell phenotypes based on spatial context ^26,27^. If contamination is not properly addressed, it may introduce systematic biases that propagate through downstream predictions, underscoring the need for caution when interpreting model outputs.

We demonstrated that incorporating snRNA-seq as a reference significantly enhances cell-type annotation. Additionally, RCTD’s doublet model provides a useful means of assessing sample quality and segmentation accuracy by quantifying the prevalence of doublets. We further leverage this model within SPLIT to decompose mixed signals and correct for transcript diffusion, thereby improving cell-type resolution. While matched references from the same sample are preferable—particularly for distinguishing malignant populations—we show that external references also perform well, underscoring the robustness and generalizability of reference-based approaches.

This flexibility enables broader adoption of RCTD-based annotation and, by extension, SPLIT, even when matched snRNA-seq data are unavailable.

We also evaluated the performance and implications of background correction and segmentation strategies, including our method, SPLIT. While many correction techniques can be applied independently of the segmentation algorithm, they often raise concerns about interpretability and the risk of over-correction—particularly when used in combination with transcript-space segmentation methods like ProSeg, which already account for diffusion. This is especially evident with approaches such as ResolVI, which can substantially reduce gene counts and, as a result, may compromise statistical power. In contrast, SPLIT is less prone to these artifacts, as it operates as a reference-based signal decomposition method rather than a direct spatial correction layer. Its outputs are inherently more interpretable, and importantly, SPLIT preserves gene detection levels more effectively than alternative methods.

Segmentation in transcript space—particularly using probabilistic methods such as ProSeg—led to notable improvements in cell-type resolution, especially when paired with targeted panels. However, these gains were more modest with the 5K panel, likely due to its lower sensitivity and the increased complexity of performing segmentation in a higher-dimensional space. Despite these challenges, transcript-based segmentation remains fully compatible with correction methods like SPLIT, allowing them to be effectively combined for enhanced resolution. Notably, applying SPLIT to the 5K panel improved cell-type separation when using the default 5 μm extension, highlighting its utility even under standard segmentation conditions.

An important consideration when using SPLIT is its potential to assign multiple phenotypes to a single cell, requiring careful downstream interpretation of primary versus secondary identities. In cases of substantial contamination—particularly in low RNA-content cells such as T cells—the true phenotype may be incorrectly classified as secondary. This underscores the need for computational strategies to more accurately interpret and resolve secondary phenotypes. In our current implementation, we retain the primary phenotype but optionally allow phenotype shifts through a simple procedure we refer to as SPLIT-shift. Future work could explore more advanced approaches, such as reassigning transcripts from contaminating profiles to neighboring cells or developing criteria to determine which phenotype most likely reflects a cell’s true identity—potentially incorporating morphological features to guide this inference. It is also important to recognize that both cell-type annotation and SPLIT rely on the assumption that all relevant cell types are present in the reference dataset. The absence of specific cell types can compromise RCTD-based annotation, potentially resulting in artificial doublet assignments as the model attempts to account for unrepresented variation. For instance, cycling tumor cells may be incorrectly decomposed into a combination of cycling T cells and non-cycling tumor cells. This represents a critical caveat that users should consider when interpreting annotation and decomposition results.

## Methods

### Human samples

All patients provided informed consent for the use of the tumor samples in this study. The protocol used for sample collection was approved by the local ethics committee (CER-VD, BASEC ID 2016-02094).

### Datasets

#### Study design

Data were generated from 10 formalin-fixed paraffin-embedded (FFPE) non-small cell lung cancer (NSCLC) samples and 17 FFPE breast cancer samples. Xenium spatial transcriptomics was performed on all samples using different gene panels, with the 5K panel specifically applied to lung samples. Additionally, matched single-nucleus RNA-seq (snRNA-seq) and multiplexed immunohistochemistry (IHC) data were generated for selected samples, as detailed below and in Figure 1.

#### Xenium

Xenium lung tissues were processed using three gene panels in parallel, the 10x Human Lung panel, a custom immuno-oncology panel (Custom IO; see Supplementary Table 2 for gene list), and the 10x Prime 5K Human Pan Tissue & Pathways panel, allowing an investigation into how the choice of a gene panel may affect downstream analyses. For breast samples, we used the 10x Human Breast panel.

#### snRNA-seq

snRNA-seq data were obtained from FFPE blocks from eight patients, four per disease group. Details about this dataset and its methodology are available in the publication by Dong et al^2^.

#### IHC

Two post-Xenium slides—one per disease—containing a total of five tissue sections were immunohistochemically stained for DAPI (Spectral DAPI, AKOYA), CD8 (Cellmarque, Clone SP16, 108R), and pan-cytokeratin (panCK, Dako, Clone AE1/AE3, IS053) in autostainer (ventana Discovery Ultra, Roche) using Tyramide Signal Amplification system (OPAL, Akoya), allowing us to identify CD8^+^ T cells and malignant cells. Antigenicity was reduced in some samples, particularly those processed with the multimodal staining protocol used for Xenium segmentation, leading to failed IHC staining in certain cases. As a result, IHC yielded usable staining quality in only five samples.

### Data preprocessing and quality control

#### snRNA-seq

Filtered barcode counts from CellRanger were analysed using the Seurat package^28^ (v. 5.0.1) in R (v. 4.3.2, https://www.R-project.org/). Cells with at least 200 detected genes and fewer than 20% of reads mapping to mitochondrial genes were retained. To recover neutrophils, the gene threshold was lowered to 100 genes per cell, and cells forming a distinct cluster with high expression of canonical neutrophil markers (i.e., FCGR3B, S100A9, IL1R2, CSF3R, FPR1, NAMPT) were retained.

Raw counts were normalized using SCTransform, and the top 3000 variable genes across samples were selected using *SelectIntegrationFeatures*. Dimensionality reduction was done using principal component analysis (PCA). Clustering was done via Seurat’s shared nearest neighbor (SNN) modularity optimization algorithm (*FindNeighbors* and *FindClusters*) using 30 principal components and resolutions between 0.4 and 0.8.

Clusters were annotated based on the top differentially expressed genes and canonical markers. Significant markers were identified with *Seurat::FindAllMarkers()* (two-sided Wilcoxon’s rank sum test, Bonferroni correction, adjusted p < 0.05, log fold change > 1). Sub-clustering enabled finer annotation, resulting in four hierarchical annotation levels. Cell type labels were standardized to match ontology naming conventions. This dataset was used as a matched reference for RCTD cell-type annotation of Xenium data.

Annotation Level 2.1 was added for visualizing cell types on Xenium. As an extension of the broader Level 2, it distinguishes CD8^+^ T cells and supports validation with IHC staining.

#### Xenium

##### 10x segmentation

To facilitate segmentation, Xenium’s standard output includes a DAPI-stained image for nuclear localization along with the x, y, and z coordinates of individual transcripts. Segmentation algorithms can leverage this information to assign transcripts to individual cells. Unless the multimodal segmentation kit is used, the default 10x segmentation algorithm expands from the nucleus outward—up to 5 µm or until it reaches the boundary of an adjacent cell.

In our study, the multimodal segmentation kit was applied only to the 5K panel. Therefore, we used the default 5 µm nuclear expansion-based segmentation for all other panels. In practice, segmentation results are identical for a substantial proportion of cells, as the multimodal method defaults to the 5 µm nuclear expansion when membrane staining is weak or absent. In either case, we consider these approaches to provide reasonable and consistent baselines for our analyses.

##### Alternative segmentations

Several alternative segmentation methods have been developed to improve upon the default 10x approach described above. In this study, we evaluated Baysor (v0.7.0, https://github.com/kharchenkolab/Baysor), ProSeg (v2, https://github.com/dcjones/proseg, commit ef2d1ca8c535fd911b7cd37da47c84de98e784a4), and Segger (a version adapted by us at https://github.com/bdsc-tds/segger_dev, commit 4bf56dec2a364de8eee4fcab663e798eb106e21a)—all of which utilize transcript-level information to guide segmentation. We applied these methods with default arguments to all Xenium samples. Specifically, when running Segger, we used 6,000 tokens for samples from the 5K panel and 500 tokens for others. These methods represent robust alternatives and serve as valuable baselines for benchmarking segmentation performance.

##### Quality control

Cells with less than 10 counts or 5 genes were filtered out, as well as genes expressed in fewer than 10 cells.

#### IHC

##### Analysis

Xenium and IHC images were co-registered in QuPath (v0.5.1) using the warpy extension (https://github.com/BIOP/qupath-extension-warpy). Xenium’s cell segmentation was loaded as a GeoJSON file, and an object classifier was trained within QuPath to classify cells based on protein expression.

### Downstream analyses

#### UMAP visualization

UMAPs were computed for log-normalized snRNA-seq and for Xenium data at the sample and panel level using default scanpy ^29^ parameters for PCA, kNN search and UMAP, except kNN n_neighbors which was set to 50.

#### Xenium annotation with RCTD

We performed cell type annotation of Xenium data using the RCTD algorithm in doublet mode, leveraging both matched snRNA-seq and external scRNA-seq references (see Data availability section). Each sample was annotated independently. To enhance the specificity of tumor cell detection, matched references were restricted to include only tumor cells derived from the corresponding donor, when donor-specific cells were available. Reference datasets were filtered to remove cells with fewer than 10 or more than 2000 unique molecular identifiers (UMIs), and cell types represented by fewer than 25 cells were excluded. RCTD was run on raw UMI count data from both the reference and Xenium data. Due to the inherently lower UMI counts in single-cell resolution Xenium data, we applied a more permissive filtering strategy: cells with fewer than 10 total UMIs were excluded, and *UMI_min_sigma* parameter was reduced to 100 to account for the reduced transcript coverage compared to spot-based spatial transcriptomics in the original RCTD application. To improve the robustness of RCTD annotation, we provided the *class_df* parameter, which groups cell types into broader classes. Finally, we applied mild post-processing to the RCTD results, removing an arbitrary second cell type in highly confident singlets—specifically, singlets that showed no evidence of a secondary signal.

#### Spatial diffusion and contamination metrics

*Neighborhood abundance of the secondary cell type (neighborhood w2)* is defined as the average weight of the secondary cell type signal within a cell’s spatial neighborhood.

*Cell-type diffusion potential* is calculated as the mean neighborhood w2 across all cells sharing the same secondary cell type.

*Cell-type transcript abundance* is defined as the product of the mean transcript count (nCount) for a given cell-type profile and its proportion within the sample.

#### SPLIT correction

To reduce contamination from secondary cell type signals in an observed expression profile, we leveraged RCTD output. RCTD provides, for each cell, the primary and secondary cell type labels along with doublet weights *w*_1_ and *w*_2_ = 1 − *w*_1_, which reflect the relative contributions of the primary and secondary reference profiles to the observed expression. Using these weights and the reference profiles, we computed a purified expression profile designed to separate the contribution of the primary cell type from the secondary cell type. Specifically, for each cell:

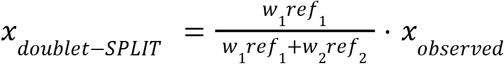

where *ref*_1_ and *ref*_2_ are the average reference profiles of the primary and secondary cell types, respectively. Intuitively, this ratio (used as a scaling factor) is an estimate of the expected fraction of transcripts coming from the primary cell type. The complement, namely 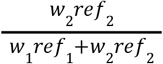, being the expected fraction of transcripts coming from the secondary cell type.

We call this approach doublet-SPLIT and by default apply it to cells classified as “doublets_certain” and “singlets” that showed signs of contamination—i.e., cells for which both a primary and secondary cell type were confidently assigned by RCTD.

For cells labeled as “doublets_uncertain”, only the primary cell type is confidently assigned, and the secondary identity is considered ambiguous. To purify these cells, we performed a full-SPLIT as:

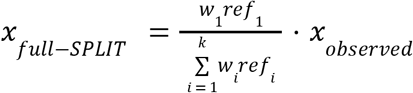

In addition, we observed that RCTD occasionally swaps the primary and secondary cell types. To account for this, we implemented a label-correction step termed SPLIT-shift. If a cell has primary label *ct*_1_ and secondary label *ct*_2_, but its transcriptomic neighborhood is primarily composed of *ct*_2_ cells, we assume a potential label swap and perform purification using *ct*_2_ as the primary identity, namely:

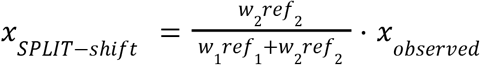

To support SPLIT-shift decisions, we also constructed a transcriptomic neighborhood for each cell, defined by a k-NN graph (k=10) based on euclidean distance in PCA space.

#### Which cells to SPLIT

We offer several modes to decide which cells should be purified:

- `Default`: applies doublet-SPLIT to singlets and doublets_certain, and full-SPLIT to doublets_uncertain.
- `balance_spot_class_based`: singlets remain unchanged; doublet-SPLIT is applied to doublets_certain, and full-SPLIT to doublets_uncertain.
- `balance_score_based`: singlets and doublets_certain undergo doublet-SPLIT only if they show evidence of contamination—specifically, if they are surrounded by cells of their assigned secondary type. Doublets_uncertain still receive full-SPLIT; all others remain unchanged. To support this decision-making, we introduce a spatial neighborhood–based metric, `neighborhood_weight_second_type` (referred to as neighborhood w2 in Figure 3), which represents the average proportion of a cell’s secondary type found within its local spatial neighborhood. In this study, we defined each cell’s spatial neighborhood using a k-nearest neighbor approach with (k=20) based on euclidean distance in physical space. To ensure locality, we removed all edges longer than 15 µm.

##### Alternative count correction

#### ResolVI^30^

is a probabilistic model that refines spatial transcriptomic data by resolving cell type mixtures using variational inference. In this study, we applied ResolVI using default parameters, with models trained either in an unsupervised manner or supervised using Level 2.1 cell-type labels.

#### Ovrlpy^24^

combines a vertical sub-slicing strategy with an unsupervised, segmentation-free analysis of spatial transcriptomics data to identify and correct for potential spatial doublets. In this study, we applied ovrlpy with default parameters, with a threshold of either 0.5 or 0.7 applied to the pixel signal integrity map, as recommended in the original preprint and through personal communication with the authors (https://github.com/HiDiHlabs/ovrl.py/issues/40) respectively.

##### Comparing segmentations and count correction methods

Computational procedures and metrics for assessing cell type separation, batch effect, data quality and potential spatial signal contamination. Analyses focused on quantifying specificity (i.e., potential contamination between neighboring cell types) and sensitivity (i.e., data preservation, similarity to chromium profiles) were adapted from Mitchel et al 2025 ^14^. These were applied to each sample, before and after count correction.

#### Assessment of Cell Type Separation and Batch Effect strength

Biological conservation of cell types and batch effect strength were assessed using metrics proposed by Luecken et al, 2022 ^31^, as implemented in the scib-metrics package (https://github.com/YosefLab/scib-metrics). Malignant cells were excluded before computing these metrics because they are often sample specific.

#### Assessment of Specificity and Potential Contamination

To evaluate potential transcript diffusion or misassignment between spatially adjacent cells of different types, we reimplemented a recently proposed approach ^14^. For each pair of distinct cell types, denoted as the target cell type *i* and the potential contaminating cell type *j*, cells annotated as type *i* were assigned a binary label: 1 for those spatially adjacent (based on centroid distance <= 15 µm) to at least one cell of type *j*, and 0 for those not adjacent to any cell of type *j*. We then trained a logistic regression model to predict, based on the gene expression profile of a cell of type *i*, whether it is adjacent to a cell of type *j*.

For each sample after QC filtering, input features were log-normalized expression levels of all genes for cells of type *i*. We used the implementation from scikit-learn^32^ with default parameters, except for *max_iter* which was increased to 500 and *class_weight* set to ‘balanced’ to account for possible imbalance of target labels.

To facilitate interpretation and comparison of feature importance, estimated coefficients for each gene were scaled by multiplying them with the standard deviation of that gene’s expression across input cells of type *i*. Model performance in predicting adjacency was assessed using the average test set precision score obtained from 5-fold spatial cross-validation. Spatial cross-validation sets were obtained using scikit-learn’s bisecting k-means on spatial coordinates of cells of type i.

To assess whether the model performs better than chance, we obtained empirical p-values by comparing to the average test set precision scores from 30 models trained on random permutations of the target variable.

Genes were ranked by the magnitude of their scaled coefficients to assess their potential association with signal contamination from cell type j within cell type i. To evaluate whether these ranked gene lists were enriched for known markers of cell type *j*, we performed enrichment analyses using two approaches. For each sample, the top *N* markers (where *N* = 10, 20, 30, 40, or 50) of type *j* were identified via Wilcoxon rank-sum tests, comparing to all other cells. Enrichment among the top 20 ranked coefficients was then assessed using either a hypergeometric test implemented in scipy ^33^ or prerank Gene Set Enrichment Analysis (GSEA) normalised enrichment score (NES) ^34,35^ implemented in GSEApy ^36^.

This analysis was run across all segmentations of each Xenium sample, both before and after applying count correction. Cell types with more than 50,’000 cells were downsampled to a maximum of 25,’000 cells from each class. For a given sample, cell types were excluded from analysis if: (1) fewer than 30 cells were available in both target classes (proximal vs. distal to cell type *j*), or (2) spatial cross-validation with 5 folds could not be constructed with both classes present.

#### Assessment of Sensitivity and Data Quality

After applying QC filters (see Methods, Data preprocessing, Xenium) to raw or corrected count matrices, standard QC metrics were computed per cell, including the total number of UMIs or reads detected, and the number of genes expressed (defined as genes with > 0 counts). Mean and median values for the number of genes expressed per cell, as well as the mean UMI count per cell, were calculated across all cells.

To assess the biological fidelity of the expression profiles, we compared them to reference chromium data. Pseudo-bulk expression profiles were generated for each identified cell type in our dataset by averaging log-normalized gene expression across all cells assigned to that type. We then calculated the cosine similarity between these pseudo-bulk profiles and corresponding cell type profiles derived from Chromium scRNA-seq. The similarity calculation was performed using all shared genes.

#### Analysis coordination

Given the raw Xenium data and the curated snRNA-seq reference, we implemented a reproducible Snakemake^37^ pipeline to coordinate the above sophisticated analysis on a high performance cluster. The number of threads and the amount of memory for each task were allocated dynamically, with a limit of 48 threads and 1TB memory, and the run time was limited to three days (72 hours). Tasks that failed due to out-of-memory errors were excluded from the results.

## Code availability

Details on reproducing our analysis and analyzing other datasets can be found on GitHub (https://github.com/bdsc-tds/xenium_analysis_pipeline). SPLIT is available as an R package and can be found on GitHub (https://github.com/bdsc-tds/SPLIT).

## Data availability

Xenium data produced as part of this study will be made available upon submission.

Raw data of our matched lung and breast cancer snRNA-seq references are publicly available from cellxgene at https://cellxgene.cziscience.com/collections/bd552f76-1f1b-43a3-b9ee-0aace57e90d6

The external lung scRNA-seq reference can be found here: https://cellxgene.cziscience.com/collections/edb893ee-4066-4128-9aec-5eb2b03f8287. We used the extended atlas, containing data from 318 patients, which accounts for over 1 million cells^38^.

The external breast scRNA-seq reference, described by Wu et al, 2021^39^, can be found here: https://www.ncbi.nlm.nih.gov/geo/query/acc.cgi?acc=GSE176078.

## Acknowledgments

While independently developing SPLIT, we discovered that the RCTD source code included a deprecated internal function that closely resembled the core concept of doublet-based decomposition. This insight served as inspiration for building a more comprehensive and scalable framework, specifically designed for large-scale, sparse single-cell spatial transcriptomics data. We thank the authors of the RCTD package—particularly Dylan Cable and Rafael Irizarry—for their valuable contributions to the field and for providing a robust statistical foundation that informed the development of SPLIT. This work was partially supported by an SNF project grant number (#10004717), imCORE Research Award from ROCHE/Genentech and a sponsored research agreement with 10X Genomics.

## References

1. Marx, V. Method of the Year: spatially resolved transcriptomics. Nat Methods 18, 9–14 (2021).

2. Dong, Y. et al. Transcriptome Analysis of Archived Tumor Tissues by Visium, GeoMx DSP, and Chromium Methods Reveals Inter- and Intra-Patient Heterogeneity. 2024.11.01.621259 Preprint at 10.1101/2024.11.01.621259 (2024).

3. Janesick, A. et al. High resolution mapping of the tumor microenvironment using integrated single-cell, spatial and in situ analysis. Nat Commun 14, 8353 (2023).

4. Chen, K. H., Boettiger, A. N., Moffitt, J. R., Wang, S. & Zhuang, X. Spatially resolved, highly multiplexed RNA profiling in single cells. Science 348, aaa6090 (2015).

5. He, S. et al. High-plex imaging of RNA and proteins at subcellular resolution in fixed tissue by spatial molecular imaging. Nat Biotechnol 40, 1794–1806 (2022).

6. Choi, H. et al. Spatial Transcriptomics Reveals Spatially Diverse Cancer-Associated Fibroblast in Lung Squamous Cell Carcinoma Linked to Tumor Progression. 2024.05.16.594592 Preprint at 10.1101/2024.05.16.594592 (2024).

7. Grande, E. et al. Spatial biomarkers of response to neoadjuvant therapy in muscle-invasive bladder cancer: the DUTRENEO trial. 2025.02.07.25321742 Preprint at 10.1101/2025.02.07.25321742 (2025).

8. Jia, G. et al. Spatial immune scoring system predicts hepatocellular carcinoma recurrence. Nature 1–11 (2025) doi:10.1038/s41586-025-08668-x.

9. Cook, D. P. et al. A Comparative Analysis of Imaging-Based Spatial Transcriptomics Platforms. Genomics (2023).

10. Ren, P. et al. Systematic Benchmarking of High-Throughput Subcellular Spatial Transcriptomics Platforms. 2024.12.23.630033 Preprint at 10.1101/2024.12.23.630033 (2024).

11. Wang, H. et al. Systematic benchmarking of imaging spatial transcriptomics platforms in FFPE tissues. 2023.12.07.570603 Preprint at 10.1101/2023.12.07.570603 (2023).

12. Du, L., Kang, J., Hou, Y.Sun, H.-X. & Zhang, B. SpotGF: Denoising spatially resolved transcriptomics data using an optimal transport-based gene filtering algorithm. cels 15, 969-981.e6 (2024).

13. Ergen, C. & Yosef, N. ResolVI - addressing noise and bias in spatial transcriptomics. 2025.01.20.634005 Preprint at 10.1101/2025.01.20.634005 (2025).

14. Mitchel, J., Gao, T., Cole, E., Petukhov, V. & Kharchenko, P. V. Impact of Segmentation Errors in Analysis of Spatial Transcriptomics Data. 2025.01.02.631135 Preprint at 10.1101/2025.01.02.631135 (2025).

15. Wu, L., Beechem, J. M. & Danaher, P. FastReseg: using transcript locations to refine image-based cell segmentation results in spatial transcriptomics. 2024.12.05.627051 Preprint at 10.1101/2024.12.05.627051 (2024).

16. Petukhov, V. et al. Cell segmentation in imaging-based spatial transcriptomics. Nat. Biotechnol. 40, 345–354 (2022).

17. Jones, D. C. et al. Cell Simulation as Cell Segmentation. 2024.04.25.591218 Preprint at 10.1101/2024.04.25.591218 (2024).

18. Heidari, E. et al. Segger: Fast and accurate cell segmentation of imaging-based spatial transcriptomics data. 2025.03.14.643160 Preprint at 10.1101/2025.03.14.643160 (2025).

19. Fu, X. et al. BIDCell: Biologically-informed self-supervised learning for segmentation of subcellular spatial transcriptomics data. Nat. Commun. 15, 509 (2024).

20. Cable, D. M. et al. Robust decomposition of cell type mixtures in spatial transcriptomics. Nat. Biotechnol. 40, 517–526 (2022).

21. Cheng, J., Jin, X., Smyth, G. K. & Chen, Y. Benchmarking cell type annotation methods for 10x Xenium spatial transcriptomics data. BMC Bioinformatics 26, 22 (2025).

22. van den Brink, S. C. et al. Single-cell sequencing reveals dissociation-induced gene expression in tissue subpopulations. Nat Methods 14, 935–936 (2017).

23. Ding, J. et al. Systematic comparison of single-cell and single-nucleus RNA-sequencing methods. Nat Biotechnol 38, 737–746 (2020).

24. Tiesmeyer, S. et al. 2D, or not 2D? Investigating vertical signal integrity of tissue slices. bioRxiv (2025) doi:10.1101/2025.01.13.632601.

25. Zhang, Y. et al. Gene panel selection for targeted spatial transcriptomics. Genome Biol. 25, 35 (2024).

26. Wen, H. et al. CellPLM: Pre-training of Cell Language Model Beyond Single Cells. bioRxiv (2023) doi:10.1101/2023.10.03.560734.

27. Schaar, A. C. et al. Nicheformer: a foundation model for single-cell and spatial omics. 2024.04.15.589472 Preprint at 10.1101/2024.04.15.589472 (2024).

28. Hao, Y. et al. Dictionary learning for integrative, multimodal and scalable single-cell analysis. Nat Biotechnol 42, 293–304 (2024).

29. Wolf, F. A., Angerer, P. & Theis, F. J. SCANPY: large-scale single-cell gene expression data analysis. Genome Biol. 19, 15 (2018).

30. Ergen, C. & Yosef, N. ResolVI - addressing noise and bias in spatial transcriptomics. bioRxiv (2025) doi:10.1101/2025.01.20.634005.

31. Luecken, M. D. et al. Benchmarking atlas-level data integration in single-cell genomics. Nat Methods 19, 41–50 (2022).

32. Pedregosa, F. et al. Scikit-learn: Machine Learning in Python. J. Mach. Learn. Res. abs/1201.0490, 2825–2830 (2011).

33. Virtanen, P. et al. SciPy 1.0: fundamental algorithms for scientific computing in Python. Nat Methods 17, 261–272 (2020).

34. Korotkevich, G. et al. Fast gene set enrichment analysis. 060012 Preprint at 10.1101/060012 (2021).

35. Gene set enrichment analysis: A knowledge-based approach for interpreting genome-wide expression profiles | PNAS. https://www.pnas.org/doi/10.1073/pnas.0506580102.

36. Fang, Z., Liu, X. & Peltz, G. GSEApy: a comprehensive package for performing gene set enrichment analysis in Python. Bioinformatics 39, btac757 (2023).

37. Mölder, F. et al. Sustainable data analysis with Snakemake. Preprint at 10.12688/f1000research.29032.2 (2021).

38. Salcher, S. et al. High-resolution single-cell atlas reveals diversity and plasticity of tissue-resident neutrophils in non-small cell lung cancer. Cancer Cell 40, 1503-1520.e8 (2022).

39. Wu, S. Z. et al. A single-cell and spatially resolved atlas of human breast cancers. Nat Genet 53, 1334–1347 (2021).

