## Supplementary Figures for "From Transcripts to Cells: Dissecting Sensitivity, Signal Contamination, and Specificity in Xenium Spatial Transcriptomics"

**Supplementary Table 1. Overview of Xenium samples.** For each Xenium sample, the table includes panel information, the default segmentation used and whether matching snRNA-seq and IHC data are available.

**Supplementary Table 2. Custom IO panel genes.**

**Supplementary Figure 1. Cell class composition across panels and UMAP visualistions of singlets. a,** Distribution of RCTD-inferred cell class proportions across panels in samples segmented using either multimodal staining (5k:10x_mm_5k) or nucleus-based segmentation (5k:10x_5um). Each point represents a sample; lines connect samples from the same donor. **b,** UMAP visualization of cells annotated as singlets, grouped by panel. Cells are colored by RCTD-assigned cell types using matched Chromium reference data.

**Supplementary Figure 2. Annotation of Xenium Data Using External References. a,** RCTD-inferred cell-class composition per Xenium sample, grouped by donor. **b,** Cell type composition for all Xenium samples using external references, grouped by donor and colored by cell type. The side color bar indicates the panel. **C,** Adjusted Rand Index (ARI) comparing RCTD annotations from matched vs. external references. Color indicates the panel; * denotes samples where tumor cells were not detected. **d,** UMAPs of all cells grouped by panel, colored by RCTD-assigned cell types using external Chromium data. **e,** Confusion matrix comparing cell type annotations from external (rows) and matched (columns) reference data.

**Supplementary Figure 3. Consistency of Xenium data across patients and technical replicates in breast samples.** Integrated UMAPs of the breast samples profiled with the targeted Breast panel, faceted by sample. Cells are colored by RCTD-inferred cell type using matched Chromium data.

**Supplementary Figure 4. Consistency of Xenium data across patients and technical replicates in lung samples.** Integrated UMAPs of the lung samples profiled with the prime 5k (a), targeted Lung (b) and targeted Custom IO (c) panels, faceted by sample. Cells are colored by RCTD-assigned cell types using matched Chromium data.

**Supplementary Figure 5. Cross-panel consistency across xenium and chromium. a,** Spearman correlation of mean gene counts across technical replicates in targeted panels. Each dot represents a gene. **b–c,** Correlation of gene expression between Custom IO and (b) Prime 5k and (c) Lung panels in matched samples.
**d–g,** Scatterplots of RCTD-inferred cell type compositions across matched samples profiled with different panels: (d) Lung vs. Prime 5k (5um nuclear segmentation); (e–f) Custom IO vs. Prime 5k (with/without multimodal segmentation); (g) Custom IO vs. Lung panel. **h,** UMAPs of matched Chromium data using only genes in the corresponding targeted panels. Solid lines indicate trend lines; dashed lines indicate identity (y = x). R denotes Spearman correlation.

**Supplementary Figure 6. Cell-type profile similarity across panels and assays.** Scatterplots compare the average expression of 194 shared genes between Chromium, Xenium Lung, and Xenium 5k per cell type for two donors—0PSV (a) and 1G73 (b)—profiled with all three technologies. Solid lines indicate trendlines; dashed lines denote identity (y = x). R denotes Spearman correlation.

**Supplementary Figure 7. Cell class composition across segmentations and local diffusion of secondary cell type between singlets and doublets. a,** RCTD-inferred cell class distributions across segmentation methods, all panels and samples. **b,** Sample-level distribution of the neighborhood weight of the secondary cell type (neighborhood w2). Significance: (***) for $p < 0.001$ (two sided t-test).

**Supplementary Figure 8. Cell-type diffusion score across panels. a,** Cell-type diffusion scores (average neighborhood w2 per secondary cell type) across Xenium samples from Breast, 5k, and Custom IO panels. Each circle represents a sample; size indicates cell-type transcript abundance (see Methods). **b,** Distribution of spearman correlation between cell-type diffusion potential (mean cell-type neighborhood w2) and cell-type transcript abundance (see Methods).

**Supplementary Figure 9. SPLIT. a,** Integrated UMAPs of SPLIT-corrected cells grouped by panel. **b,** Schematic of the SPLIT-shift procedure. SPLIT decomposes mixed transcriptomic profiles into purified primary and secondary components using reference profiles and assigned weights. By default, the primary profile is retained for downstream analysis. In SPLIT-shift, if there is evidence of a primary–secondary label swap—assessed based on the transcriptomic neighborhood being composed predominantly of the secondary cell type—the secondary profile is retained instead.

**Supplementary Figure 10. Doublets and specific phenotypes. a,** IHC and morphology image highlighting cells identified as malignant (primary) with cycling lymphocytes (secondary). Cells are indicated by white arrows; contour color reflects primary RCTD annotation. Dots represent cell type–specific transcripts (tumor: brown; cycling lymphocytes: cyan). **b–e,** UMAPs of the same Xenium sample: (b) computed on raw data; (c) subset to doublets; (d) computed on data purified using the default SPLIT; (e) computed on data purified with the neighborhood-aware SPLIT. All maps are colored by the primary cell type (first row), secondary cell type (second row), cell class (third row), local diffusion of the secondary cell type (neighborhood w2) (fourth row), proliferation signature (fifth row), and purification status (sixth row).

| Supplementary Table 1. Overview of Xenium samples | | | | | | | | |
|---|---|---|---|---|---|---|---|---|
| sample | donor | tissue | disease | panel | repeat | default segmentation | matching snRNA-seq | IHC post-Xenium |
| 1G73 | 1G73 | lung | NSCLC | 5K | no | 10x MM | yes | no |
| 1GAC | 1GAC | lung | NSCLC | 5K | no | 10x MM | no | no |
| 1GDD | 1GDD | lung | NSCLC | 5K | no | 10x MM | no | no |
| 1GQ9 | 1GQ9 | lung | NSCLC | 5K | no | 10x MM | no | no |
| 1GVD | 1GVD | lung | NSCLC | 5K | no | 10x MM | no | no |
| 0PSV | 0PSV | lung | NSCLC | 5K | no | 10x MM | yes | no |
| 0PSV | 0PSV | lung | NSCLC | Lung | no | 10x 5µm | yes | yes |
| 1G73 | 1G73 | lung | NSCLC | Lung | no | 10x 5µm | yes | yes |
| 0WMU_1 | 0WMU | lung | NSCLC | Lung | yes | 10x 5µm | yes | yes |
| 0WMU_2 | 0WMU | lung | NSCLC | Lung | yes | 10x 5µm | yes | no |
| 1GA2 | 1GA2 | lung | NSCLC | Lung | no | 10x 5µm | yes | no |
| 1GAC | 1GAC | lung | NSCLC | Lung | no | 10x 5µm | no | no |
| 1GFF | 1GFF | lung | NSCLC | Lung | no | 10x 5µm | no | no |
| 1GQ9 | 1GQ9 | lung | NSCLC | Lung | no | 10x 5µm | no | no |
| 0S8R | 0S8R | lung | NSCLC | Lung | no | 10x 5µm | no | no |
| 1GDD | 1GDD | lung | NSCLC | Lung | no | 10x 5µm | no | no |
| 1GVD | 1GVD | lung | NSCLC | Lung | no | 10x 5µm | no | no |
| 0PSV_1 | 0PSV | lung | NSCLC | Custom IO | yes | 10x 5µm | yes | no |
| 0PSV_2 | 0PSV | lung | NSCLC | Custom IO | yes | 10x 5µm | yes | no |
| 1G73 | 1G73 | lung | NSCLC | Custom IO | no | 10x 5µm | yes | no |
| 0WMU | 0WMU | lung | NSCLC | Custom IO | no | 10x 5µm | yes | no |
| 1GA2 | 1GA2 | lung | NSCLC | Custom IO | no | 10x 5µm | yes | no |
| 0OE1 | 0OE1 | breast | breast cancer | Breast | no | 10x 5µm | no | no |
| 0VRA | 0VRA | breast | breast cancer | Breast | no | 10x 5µm | no | no |
| 0PD6 | 0PD6 | breast | breast cancer | Breast | no | 10x 5µm | no | no |
| 19TB | 19TB | breast | breast cancer | Breast | no | 10x 5µm | no | no |
| 1BCA | 1BCA | breast | breast cancer | Breast | no | 10x 5µm | no | no |
| 1FO7 | 1FO7 | breast | breast cancer | Breast | no | 10x 5µm | no | no |
| 1GUC | 1GUC | breast | breast cancer | Breast | no | 10x 5µm | no | no |
| 1GV1 | 1GV1 | breast | breast cancer | Breast | no | 10x 5µm | no | no |
| 1H27 | 1H27 | breast | breast cancer | Breast | no | 10x 5µm | no | no |
| OPHI_1 | OPHI | breast | breast cancer | Breast | yes | 10x 5µm | no | no |
| OPHI_2 | OPHI | breast | breast cancer | Breast | yes | 10x 5µm | yes | yes |
| OPHI_3 | OPHI | breast | breast cancer | Breast | yes | 10x 5µm | yes | no |
| 1256 | 1256 | breast | breast cancer | Breast | no | 10x 5µm | yes | no |
| 1GVR | 1GVR | breast | breast cancer | Breast | no | 10x 5µm | yes | no |
| 1FHZ | 1FHZ | breast | breast cancer | Breast | no | 10x 5µm | yes | yes |
| 1ESQ | 1ESQ | breast | breast cancer | Breast | no | 10x 5µm | no | no |
| 14LZ | 14LZ | breast | breast cancer | Breast | no | 10x 5µm | no | no |
| 0U9A | 0U9A | breast | breast cancer | Breast | no | 10x 5µm | no | no |
| 1FYB | 1FYB | breast | breast cancer | Breast | no | 10x 5µm | no | no |

| **Supplementary Table 2. Custom IO panel genes.** | | | | | | | |
|---|---|---|---|---|---|---|---|
| gene | | | | | | | |
| ACTA2 | CD40 | CXCL8 | HLA-A | INMT | NCAM1 | SOX2 | WT1 |
| ADAM12 | CD40LG | CXCL9 | HLA-B | INSM1 | NCR3 | SOX9 | XCL1 |
| ADAMDEC1 | CD44 | CXCR1 | HLA-C | ISG15 | NEUROD1 | SPARC | XCL2 |
| ADH7 | CD5 | CXCR2 | HLA-DPA1 | JAML | NKG7 | SPP1 | ZNF683 |
| APOE | CD52 | CXCR3 | HLA-DQA1 | JCHAIN | NOX1 | STAT1 | |
| AR | CD63 | CXCR4 | HLA-DQA2 | JUNB | NPNT | STAT2 | |
| AREG | CD74 | CXCR5 | HLA-DRA | KDR | NR3C1 | STAT3 | |
| ARG1 | CD79A | CXCR6 | HLA-DRB5 | KIT | NR4A2 | STAT4 | |
| ASAH1 | CD79B | DERL3 | HLA-E | KLF4 | NUSAP1 | SYP | |
| ASCL1 | CD80 | DES | HMGB1 | KLRB1 | PADI4 | TACSTD2 | |
| ATG16L2 | CD82 | DNASE1L3 | HSPA1A | KLRC2 | PBXIP1 | TAGLN | |
| ATP5MC2 | CD8A | DUSP4 | HSPA6 | KLRD1 | PCNA | TAP1 | |
| AXL | CD8B | EGF | HSPH1 | KLRF1 | PDCD1 | TAP2 | |
| BANK1 | CDH1 | EGFR | ICOS | KLRG1 | PDGFRA | TBC1D4 | |
| BATF | CDH11 | EPCAM | ICOSLG | KRT14 | PDGFRB | TCF7 | |
| C1QA | CDH2 | ERBB2 | IDO1 | KRT17 | PECAM1 | TCL1A | |
| C1QB | CDK1 | ERBB3 | IFI27 | KRT20 | PI16 | THY1 | |
| C1QC | CEACAM1 | EREG | IFI30 | KRT5 | PI3 | TIGIT | |
| C3 | CEACAM8 | FAP | IFI6 | KRT6A | PLA2G2A | TIMP2 | |
| CALB2 | CFP | FAS | IFIT1 | KRT7 | PMEL | TNFRSF13B | |
| CCL17 | CHGA | FASLG | IFIT2 | KRT74 | POSTN | TNFRSF18 | |
| CCL19 | CKS2 | FCER1A | IFIT3 | KRT8 | POU2F3 | TNFRSF9 | |
| CCL2 | CLCA1 | FCER1G | IFITM1 | LAG3 | POU5F1 | TNFSF13 | |
| CCL22 | CLEC10A | FCER2 | IFITM3 | LAMP3 | PRC1 | TNFSF13B | |
| CCL3 | CLEC4C | FCGR3A | IGF1 | LEF1 | PRF1 | TNFSF18 | |
| CCL4 | CLEC9A | FGFBP2 | IGF1R | LGALS2 | PROX1 | TNFSF4 | |
| CCL5 | COL19A1 | FGL1 | IGFBP7 | LGALS3 | PSMB8 | TNFSF9 | |
| CCR1 | COL1A1 | FGR | IGHG1 | LILRA4 | PSMB9 | TOP2A | |
| CCR2 | COL3A1 | FKBP11 | IGHG2 | LILRA5 | PTGDR | TOX | |
| CCR3 | CPA3 | FMR1 | IGHM | LRRC15 | RBP5 | TP63 | |
| CCR4 | CRTAM | FN1 | IKZF2 | LTB | RGS5 | TPD52 | |
| CCR5 | CSF1 | FNIP2 | IL12B | LUM | RHOH | TPSAB1 | |
| CCR7 | CSF1R | FOLH1 | IL15 | LYVE1 | RNASE1 | TRAV1-2 | |
| CD14 | CSF2 | FOXC2 | IL15RA | LYZ | RPN2 | TRDC | |
| CD19 | CSF3 | FOXJ1 | IL1B | MCAM | RSAD2 | TRDV1 | |
| CD1A | CSF3R | FOXP3 | IL1R1 | MEF2B | RTKN2 | TRDV2 | |
| CD1C | CST7 | FSCN1 | IL1RN | MET | S100A8 | TREM2 | |
| CD200 | CSTB | GAS6 | IL2 | MFAP4 | S100A9 | TRGC2 | |
| CD207 | CTLA4 | GAST | IL2RA | MITF | SALL4 | TRGV4 | |
| CD247 | CTNNB1 | GBP1 | IL2RG | MKI67 | SECTM1 | TRGV9 | |
| CD274 | CTSG | GNLY | IL3 | MME | SELENOP | TTF1 | |
| CD28 | CTSW | GPR183 | IL32 | MMP9 | SELL | TWIST1 | |
| CD300A | CX3CR1 | GZMA | IL3RA | MRC1 | SFTPA1 | TYROBP | |
| CD38 | CXCL10 | GZMH | IL4 | MS4A1 | SLC18A2 | USP18 | |
| CD3D | CXCL12 | GZMK | IL6 | MX1 | SLC40A1 | VEGFA | |
| CD3E | CXCL13 | HAVCR2 | IL7 | MYH11 | SLC4A10 | VIM | |
| CD3G | CXCL16 | HGF | IL7R | NANOG | SNAI1 | VSIR | |
| CD4 | CXCL3 | HIGD1B | IL9 | NAPSA | SOX10 | VWF | |

**Supplementary Figure 1**

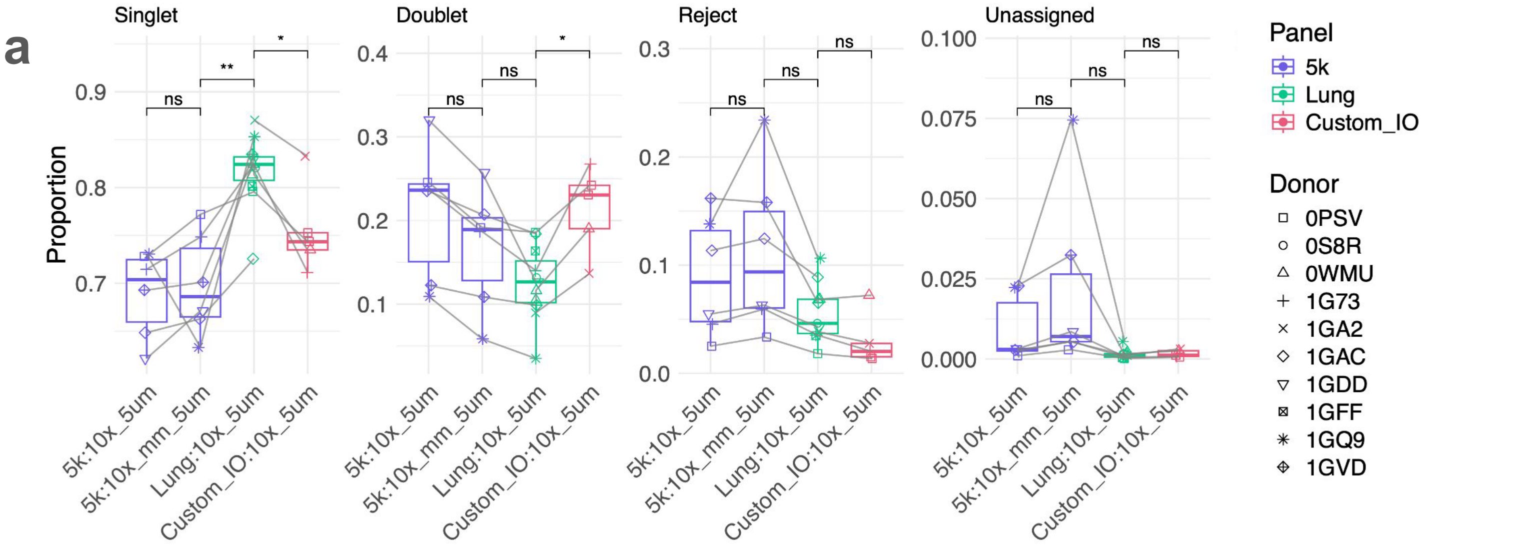


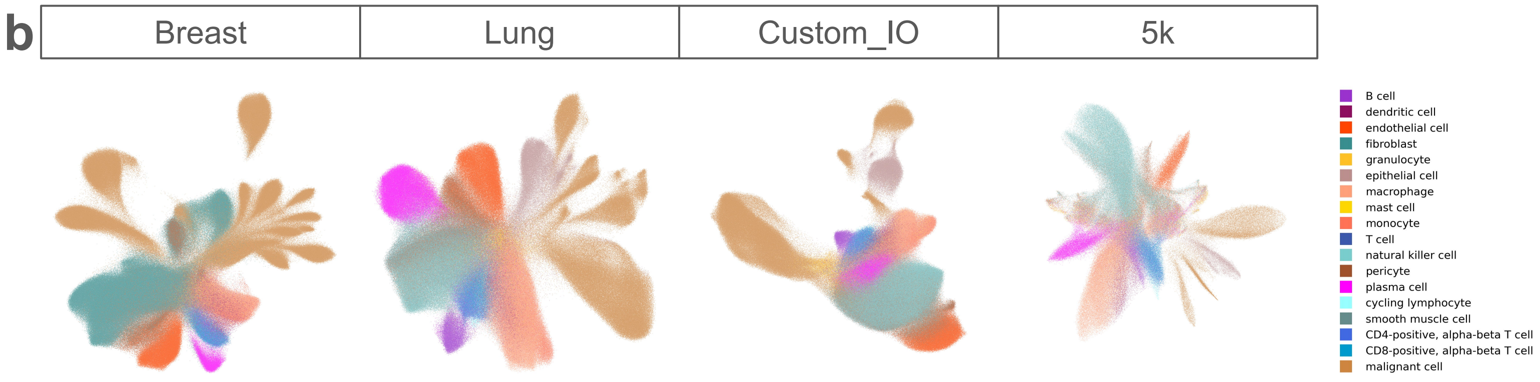

#### Supplementary Figure 2

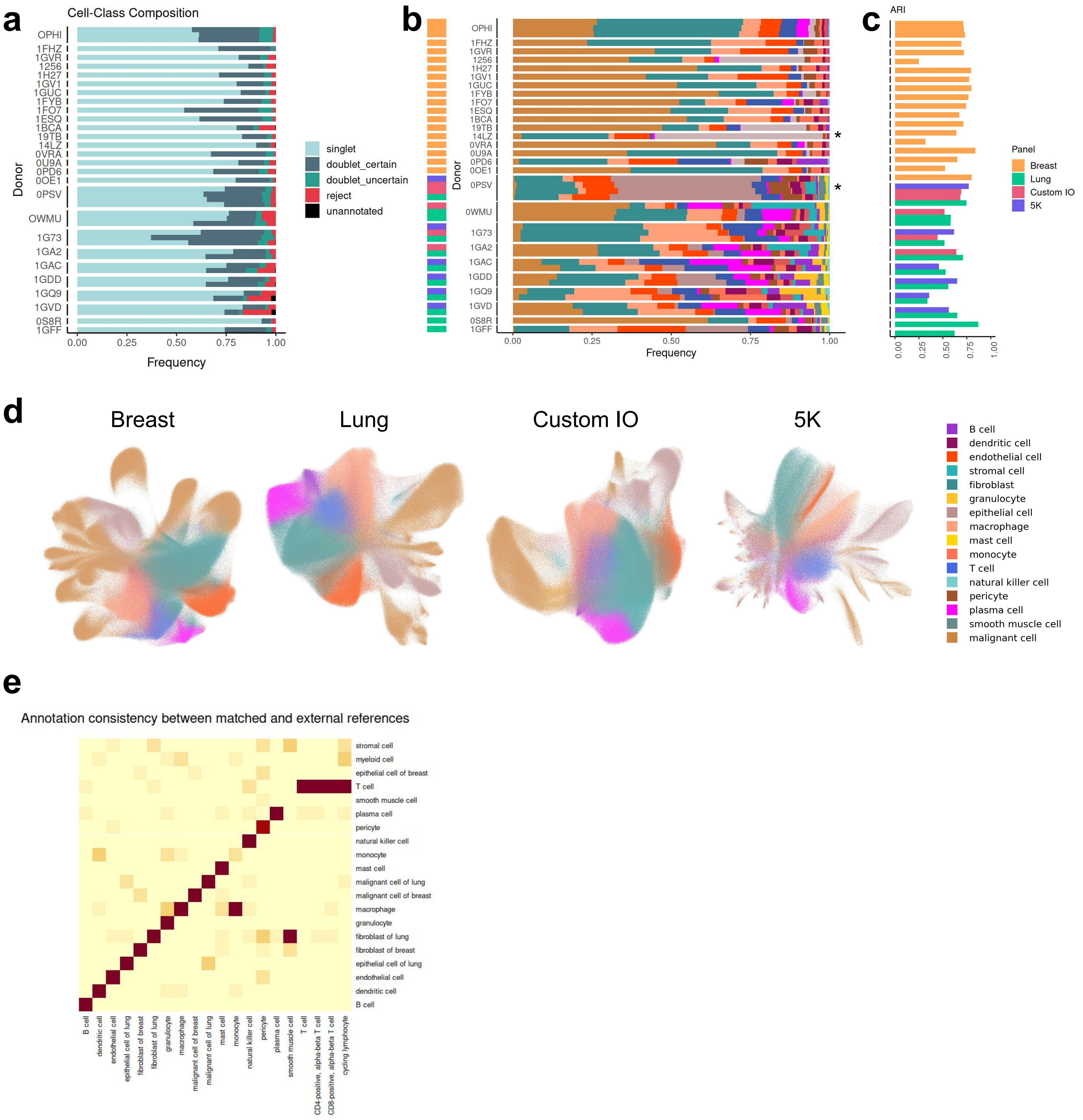

### Supplementary Figure 3

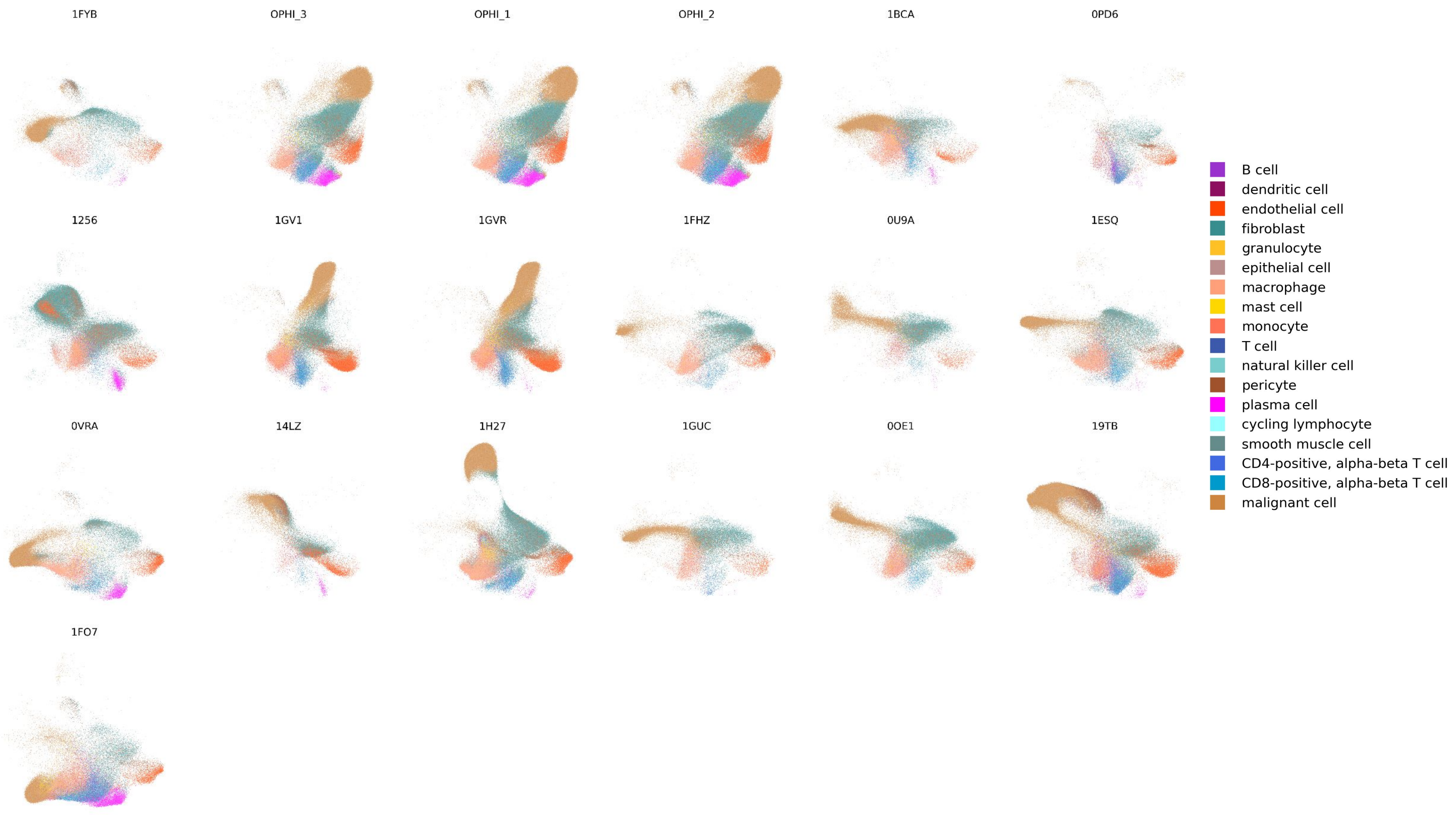

**Supplementary Figure 4**

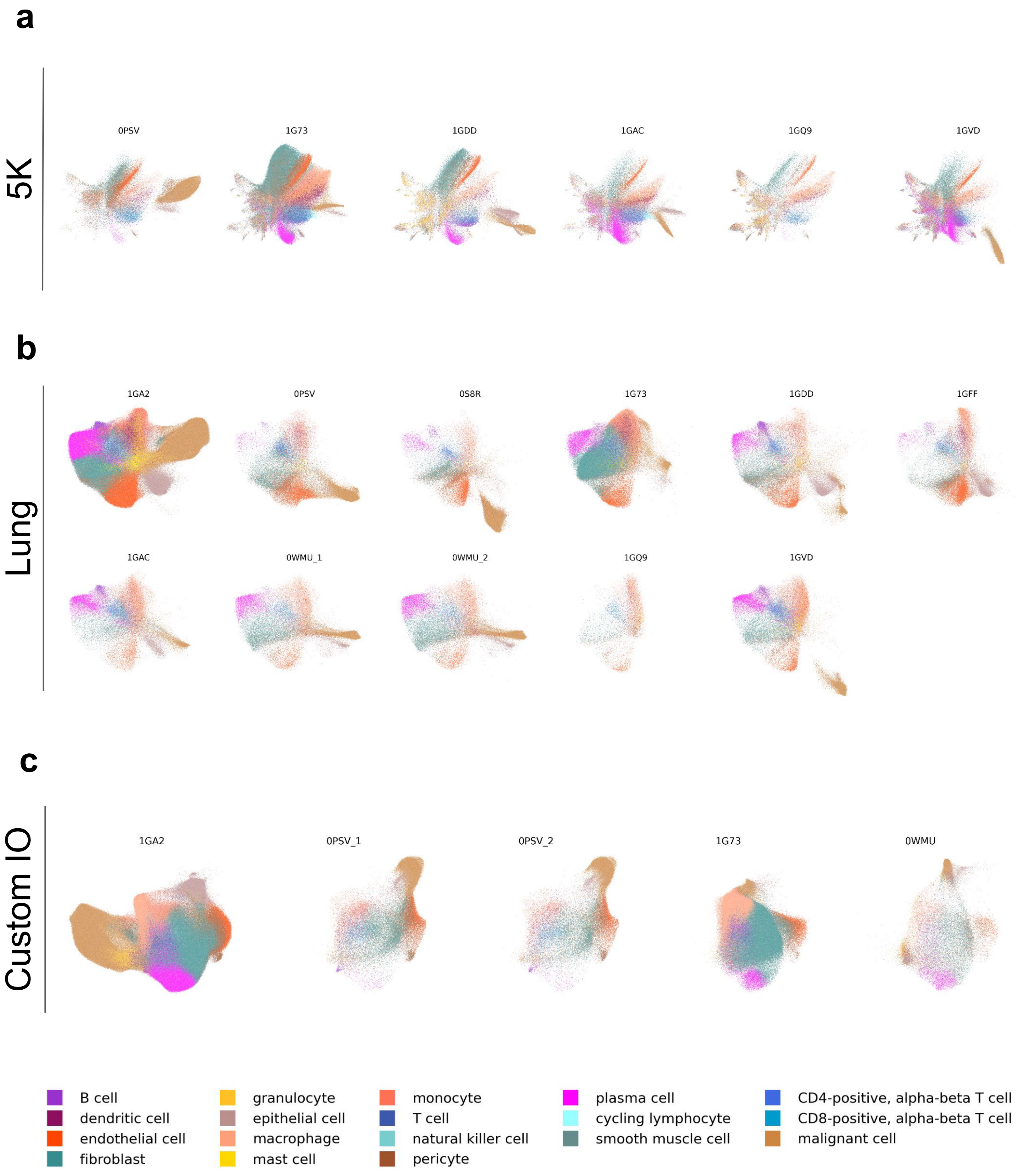

#### Supplementary Figure 5

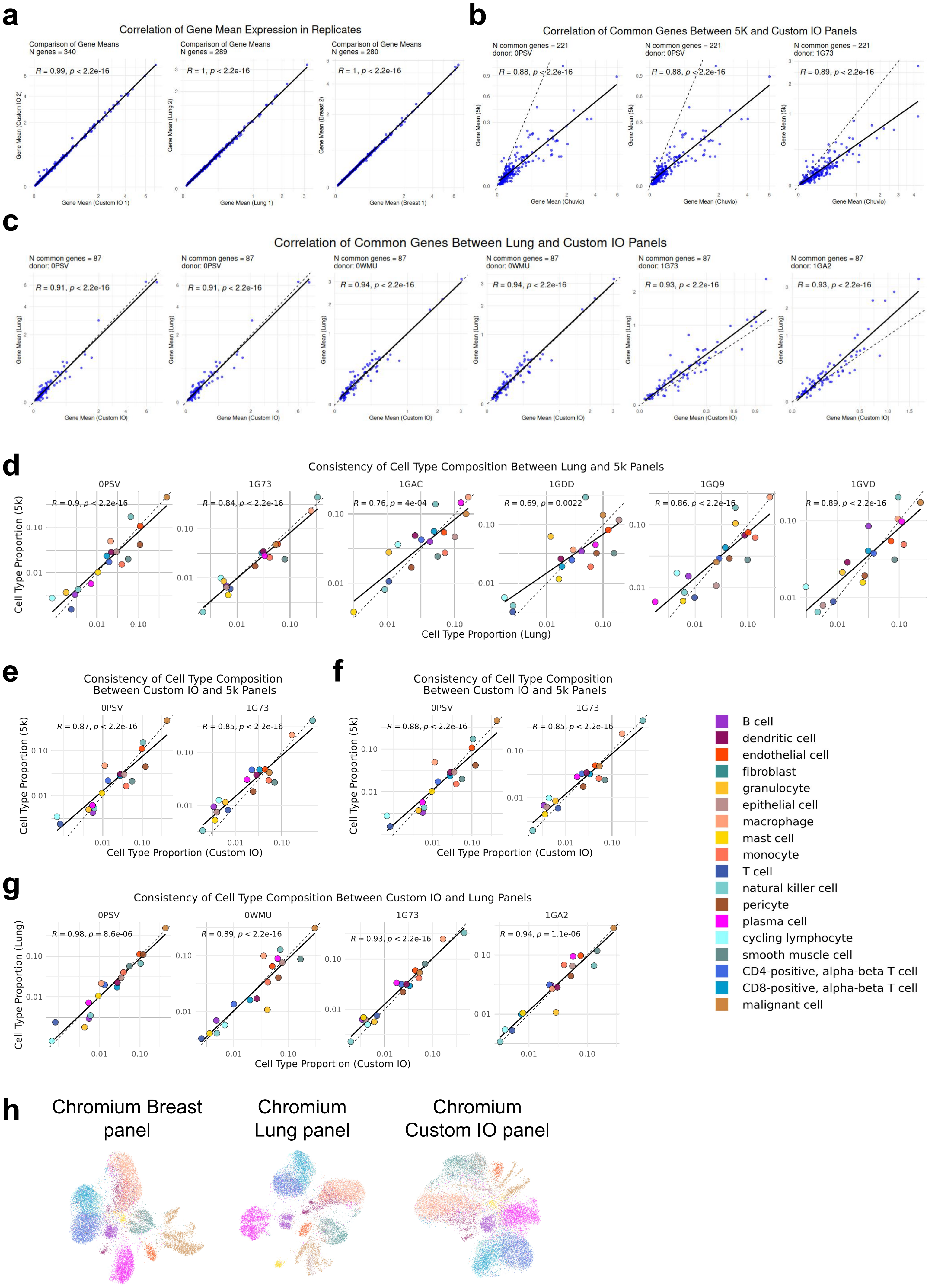

**Supplementary Figure 6**

**a**

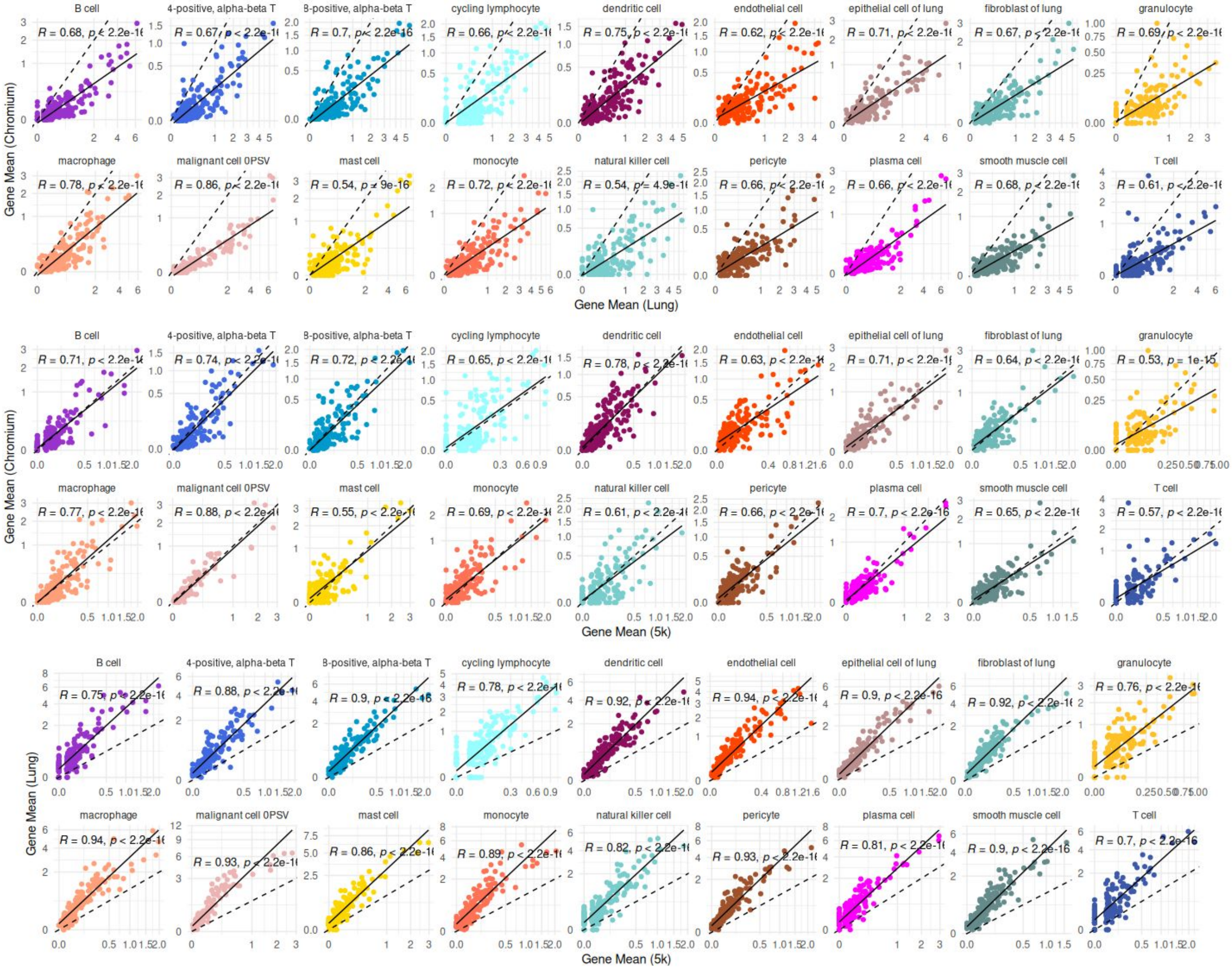


**b**

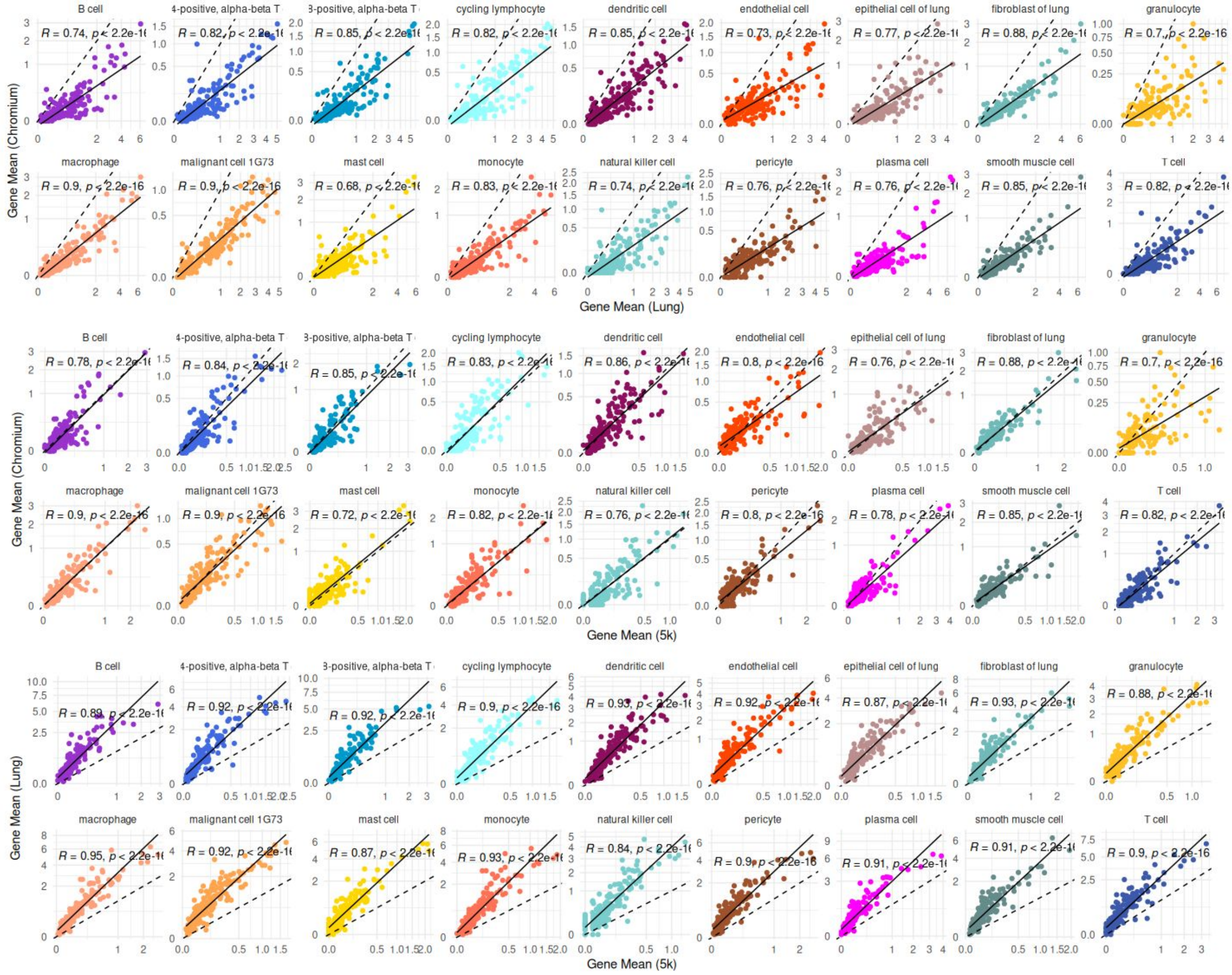

**Supplementary Figure 7**

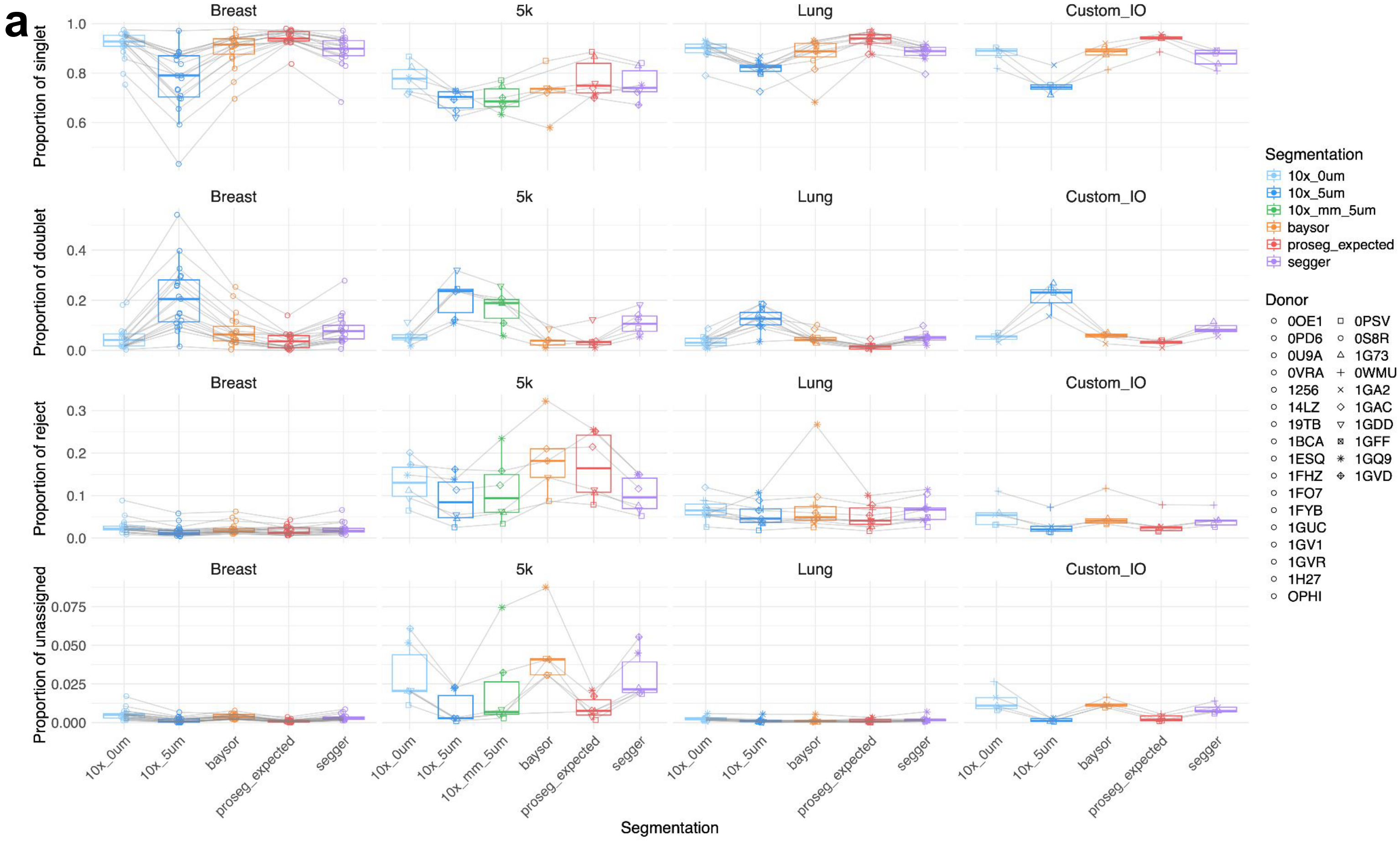


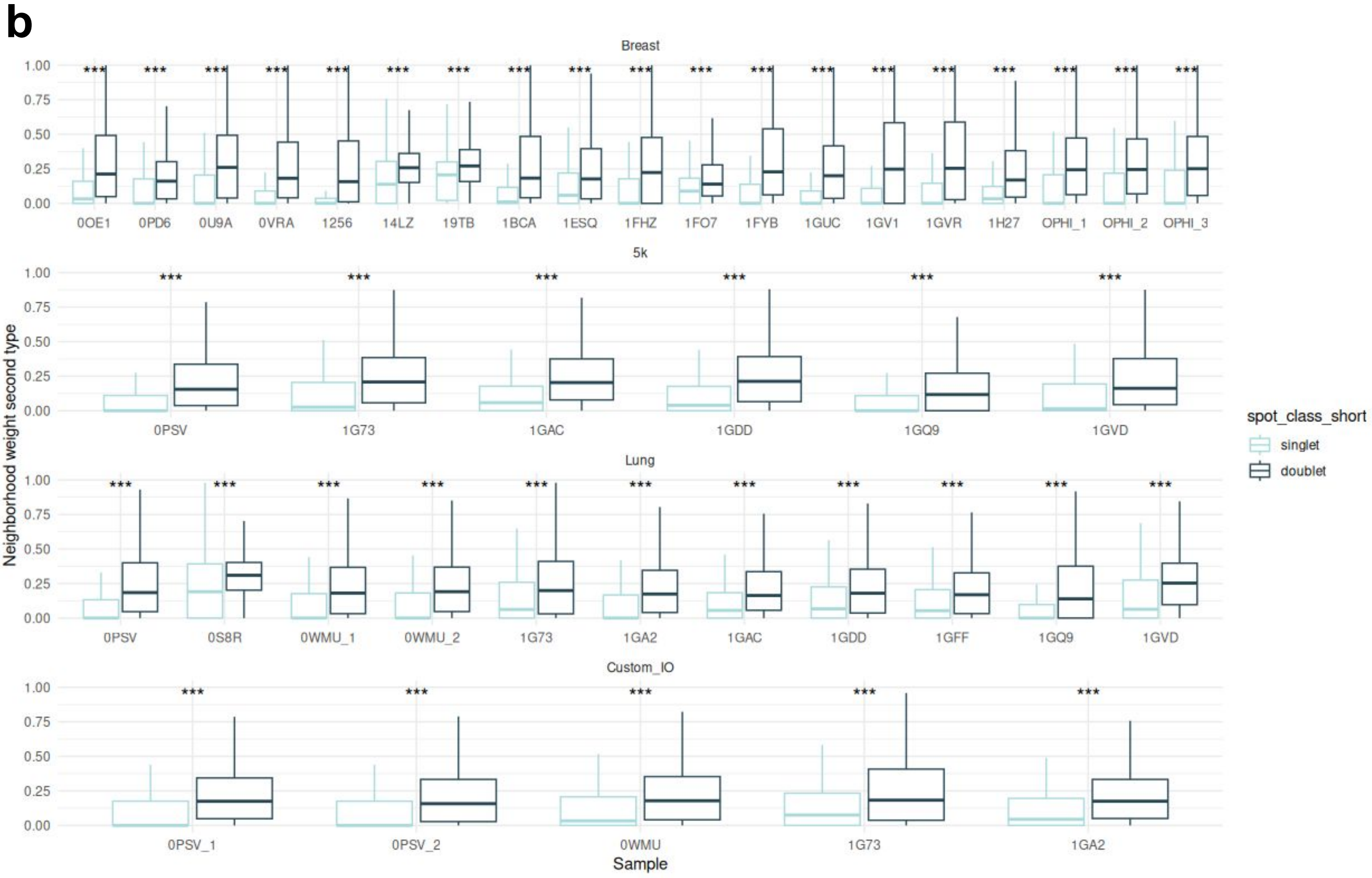

#### Supplementary Figure 8

**a**

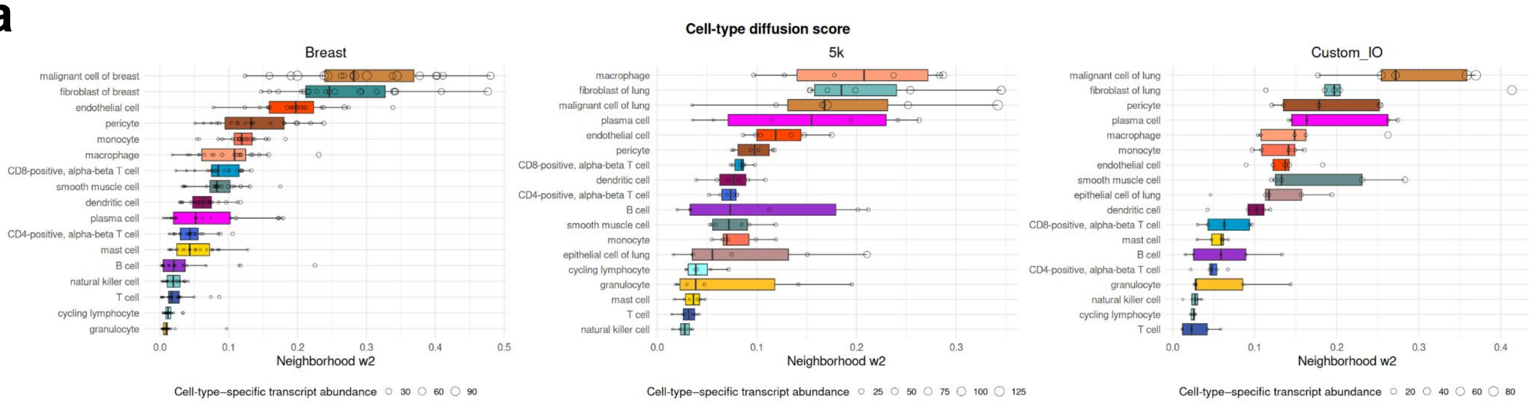


**b**

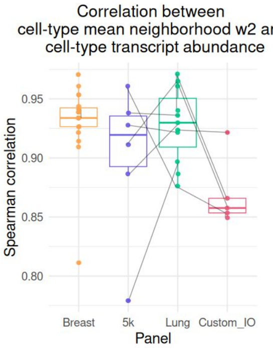

**Supplementary Figure 9**

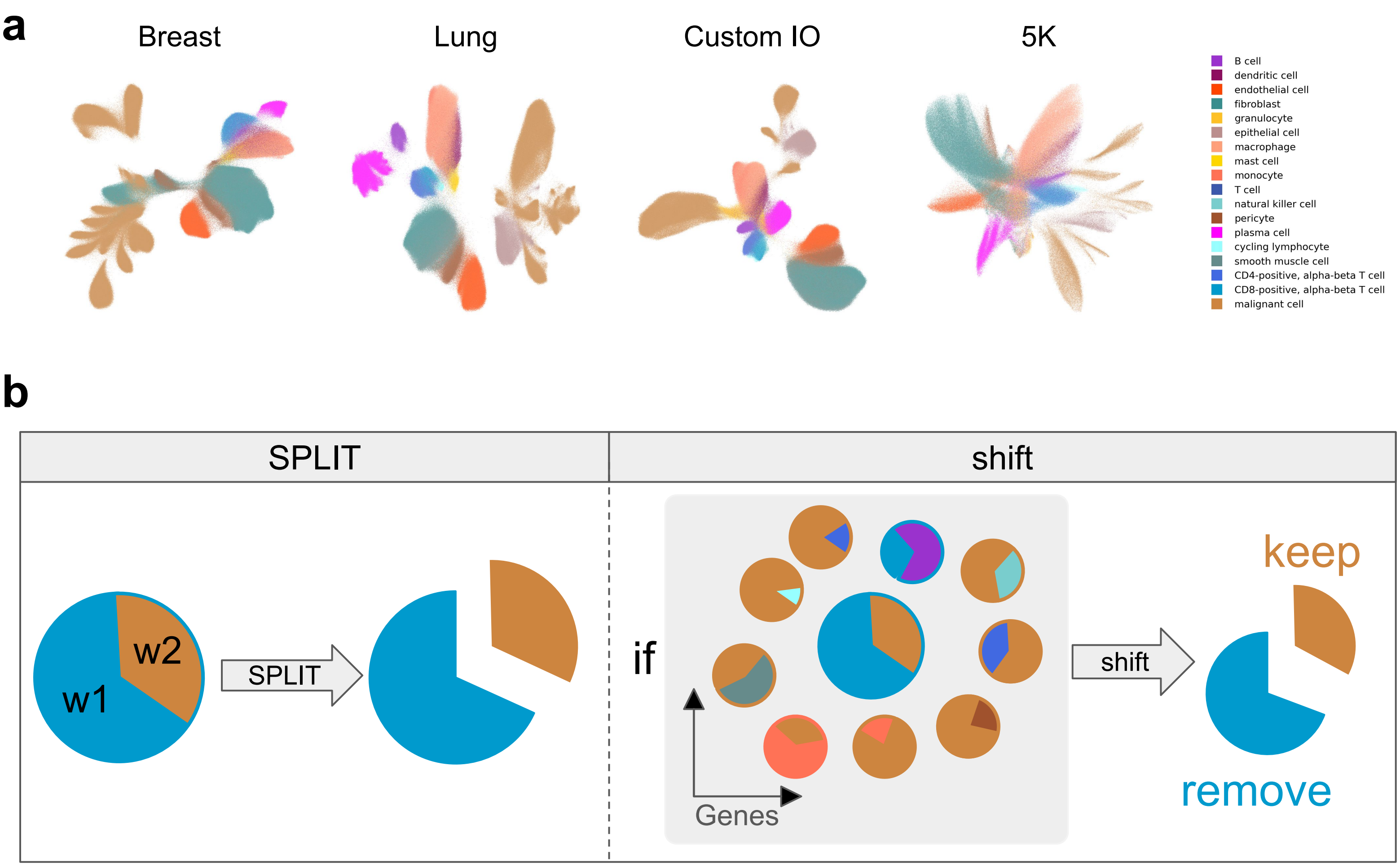

#### Supplementary Figure 10

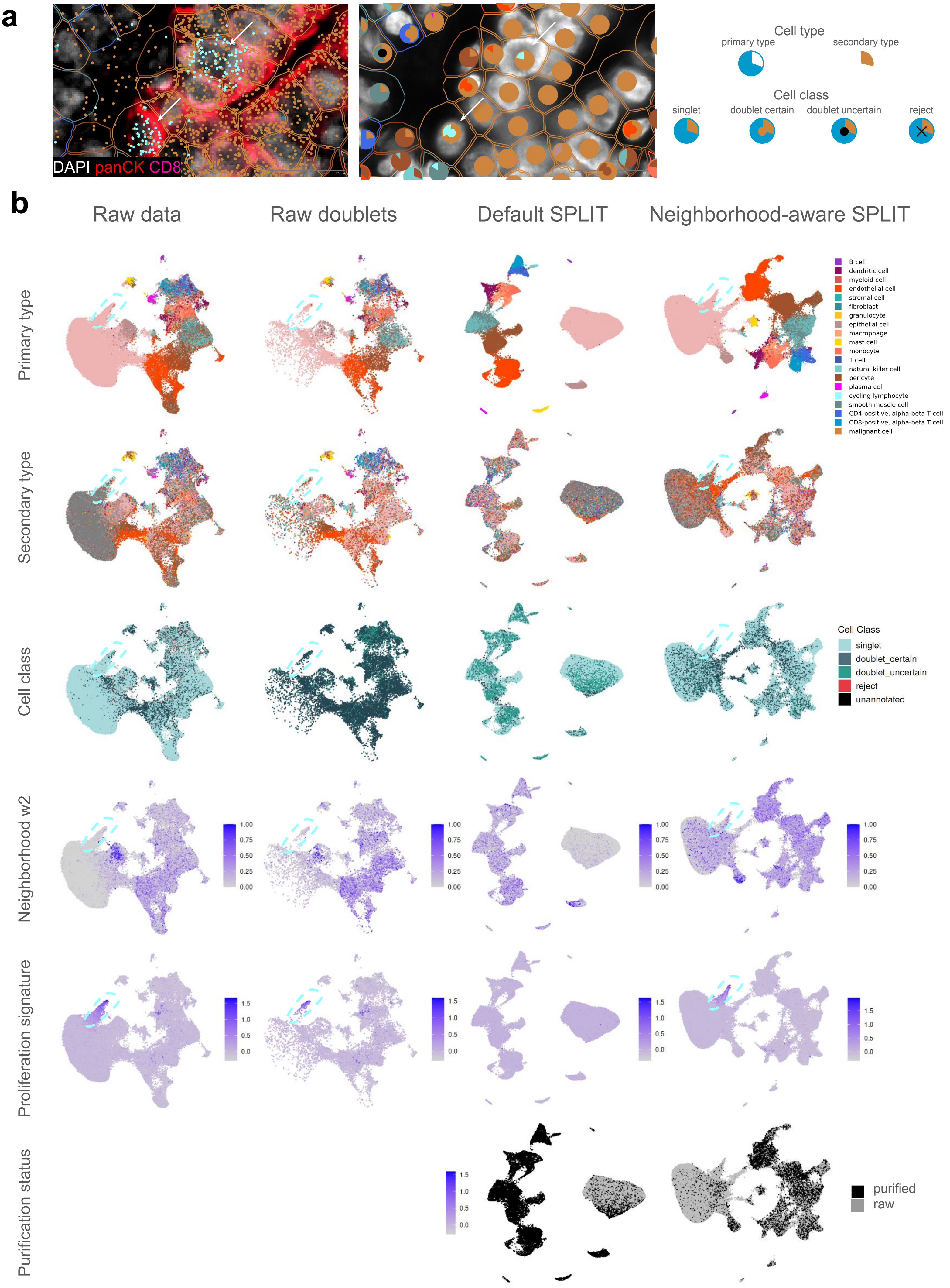
